# Mechanisms of the *FTO* locus association with obesity: Irx3 controls a sumoylation-dependent switch between adipogenesis and osteogenesis

**DOI:** 10.1101/2023.10.17.562662

**Authors:** Jan-Inge Bjune, Samantha Laber, Laurence Lawrence-Archer, Xu Zhao, Shuntaro Yamada, Niyaz Al-Sharabi, Kamal Mustafa, Pål R. Njølstad, Melina Claussnitzer, Roger D. Cox, Pierre Chymkowitch, Gunnar Mellgren, Simon N. Dankel

## Abstract

**Background:** IRX3 is implicated in genetic predisposition to obesity via the *FTO* variant locus. *IRX3* shows *FTO* risk allele-dependent upregulation specifically during early adipogenesis, leading to a shift from energy-dissipation to fat storage in mature adipocytes. However, how changes in *IRX3* expression at one developmental stage affect cellular phenotype at a later stage remains unclear. We here hypothesize that IRX3 regulates adipocyte development via transcriptional modulation of epigenetic reprogramming factors.

**Methods:** We combined ChIP-, ATAC- and RNA-sequencing to map direct Irx3 target genes in regions of open chromatin during early adipogenesis of wild-type and *Irx3-*KO preadipocytes. Gene ontology analyses was performed to identify significantly enriched biological pathways. Denaturing western blotting was used to assess sumoylation levels, and the inhibitor ML-792 was used to specifically block sumoylation. Luciferase assays were performed to estimate effects of ML-792 on Pparγ activity. Bodipy lipid staining, immunofluorescence and qPCR were employed to assess adipogenic differentiation in 3D culture. Alkaline phosphatase and Alizarine Red S staining, as well as immunofluorescence and qPCR were used to assess osteogenic differentiation in 3D culture.

**Results:** We identified more than 300 Irx3 binding sites in preadipocytes, and these were almost exclusively restricted to promoter regions, with a strong enrichment of genes related to sumoylation, histone modifications and chromatin remodeling. Genes from every step of the sumoylation cycle were bound by Irx3 and differentially expressed in response to *Irx3*-KO, leading to increased global sumoylation levels in the KO cells. Irx3 ablation and elevated sumoylation inhibited Pparγ activity and adipogenic differentiation in preadipocytes, both of which could be restored by pharmacological inhibition of sumoylation. The *Irx3-*KO cells demonstrated reduced epigenetic suppression against osteogenesis, resulting in increased osteogenesis in 3D culture. Finally, osteogenesis induced by Irx3 ablation could partially be reversed by inhibition of sumoylation.

**Conclusions:** Our study has uncovered IRX3 as a novel upstream regulator of sumoylation, and a potent controller of epigenetic regulators, both directly and indirectly via suppressing global sumoylation levels. This study indicates that the *FTO* locus promotes obesity via IRX3-mediated suppression of sumoylation, which promotes adipogenic commitment and differentiation through epigenetic programming.

**Graphical abstract:** 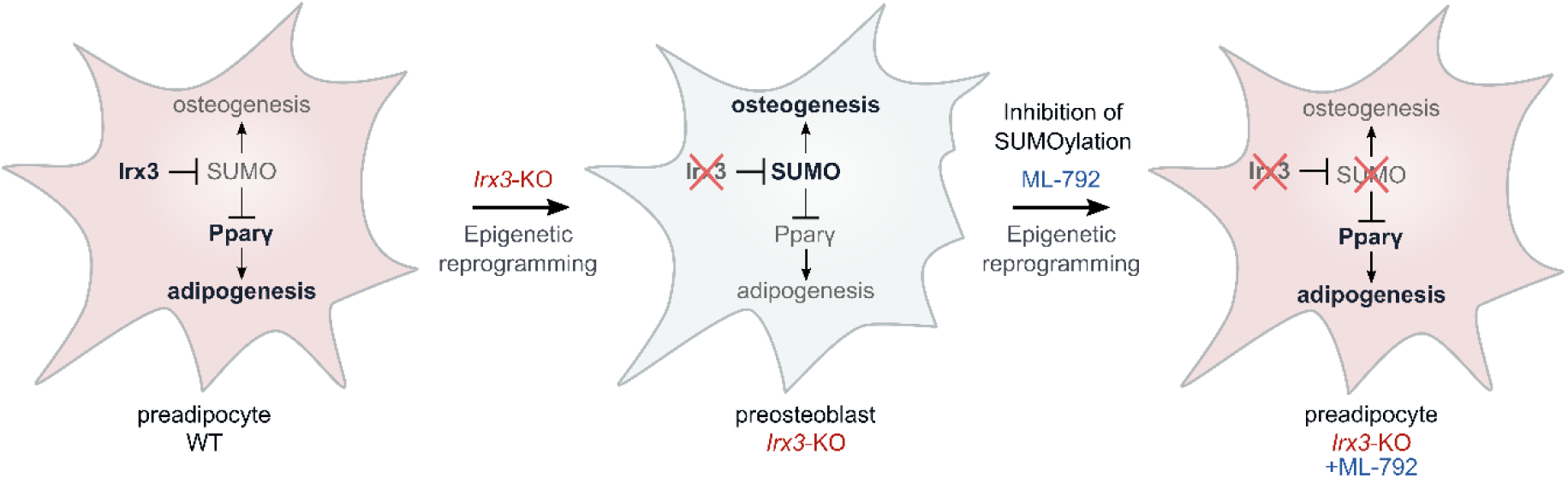

## Introduction

Common obesity has a strong genetic component that accounts for 40-80% of observed variations in body-mass index (BMI) [1,2]. Among more than 1,100 independent loci known today to be associated with BMI [3], the *FTO* locus was the first to be discovered [4–6] and has since consistently shown the strongest association with BMI across ages and ethnicities [7]. Individuals homozygous for the risk haplotype have an 1.7 increased odds of developing obesity and gain on average 3 kg more fat than homozygous protective carriers [4]. For a single, common obesity-associated variant locus, this comprises a huge effect size [3]. However, despite the impact of the *FTO* locus, the underlying mechanisms have long been debated as the associated variants are located in the introns of the gene, thereby lacking a clear function [3,8,9]. While early analyses suggested *FTO* itself could be the target gene (as reviewed in [8]), subsequent analyses by us and others showed that the haploblock is situated in a super-enhancer that regulates expression of the neighboring gene IRX3 through a 500 kb long chromatin loop [10–13].

Mechanistically, the variant-harboring enhancer was found to act on IRX3 specifically in adipose-derived mesenchymal stem cells, and only during a short time window during early stages of adipogenesis, in which the risk variant promotes a transient increase in IRX3 expression [12]. Intriguingly, this brief elevation in IRX3 levels in adipocyte precursor cells was sufficient to cause reduced beiging and diminished thermogenesis in the mature adipocytes [12]. Moreover, we recently found that complete knock-out of Irx3 in mouse embryonic fibroblasts resulted in loss of preadipocyte identity, inhibition of adipogenesis and activation of chondrogenesis [14]. These data suggest that IRX3 may have an epigenetic effect on adipocyte identity, adipogenesis and/or adipocyte function, although this has never been investigated. The IRX3 protein does not possess known chromatin remodeling domains. However, IRX3 is a homeobox transcription factor involved in a range of developmental processes [15,16], so we reasoned that it may act upstream of chromatin remodeling or histone modifying enzymes to control their expression. However, very few *direct* Irx3 target genes have been described, and to our knowledge, none related to epigenetic regulation.

The epigenetic landscape and resulting gene expression profile of any given cell largely determines the identity of that particular cell [17]. Well studied examples include the generation of induced pluripotent stem cells (iPSCs) from adult fibroblasts [18] and *vice versa;* the commitment and differentiation of pluripotent stem cells into specialized somatic cell types [19,20]. Changes in the epigenetic status, and thereby cell identity, is tightly controlled by an array of histone modifying enzymes [21,22] and chromatin remodellers [19], whose activity and function in turn can be modulated by post-translational modifications (PTMs) [23].

One PTM that is particularly involved in modulation of chromatin remodeling, epigenetic regulation and transcriptional activity is sumoylation [24–26]. SUMO (Small Ubiquitin-like Modifier) consists of three main members; Sumo-1, Sumo-2 and Sumo-3. The two latter share 97% sequence identity and are often referred to as Sumo-2/3 [27]. All SUMO isoforms are expressed as immature prepeptides that require enzymatic maturation followed by a series of enzymatic transfer steps before being conjugated to the target proteins (reviewed in [27]. While these enzymes, which collectively are termed the SUMO machinery, have been described, and the critical importance of SUMO in determining cellular identities such as discriminating totipotent versus pluripotent stem cells [28,29] is well established, the upstream regulation of this machinery as well as of SUMO itself is completely unknown.

In this study, we aimed to gain new insight into the molecular mechanisms underlying the obesity-associated enhancer in the *FTO* locus by uncovering genes that are under direct transcriptional control of Irx3 in developing preadipocytes. We identified both *Sumo* and most of the sumoylation machinery to be directly controlled by Irx3, and show how sumoylation mediates an Irx3-dependent switch between adipogenic and osteogenic cell fate.

## Results

### Irx3 binds to the promoters of the SUMO machinery and chromatin remodelers

To identify direct target genes of Irx3 in developing adipocytes, we performed ChIP-seq experiments in preadipocytes isolated from mouse inguinal white adipose tissue (iWAT) one day before (day -1) and one day after (day 1) initiation of differentiation, as well as from gonadal WAT (gWAT) on day 1 of differentiation (Figure 1A). We applied a stringent filter in which only peaks with q-value below 10e-4 and enrichment above 10-fold were considered. We identified 310 high-confidence peaks across both depots and timepoints, revealing a dramatic increase in Irx3 binding events following initiation of differentiation (Figure 1A and Supplementary file 1).

**Figure 1:**
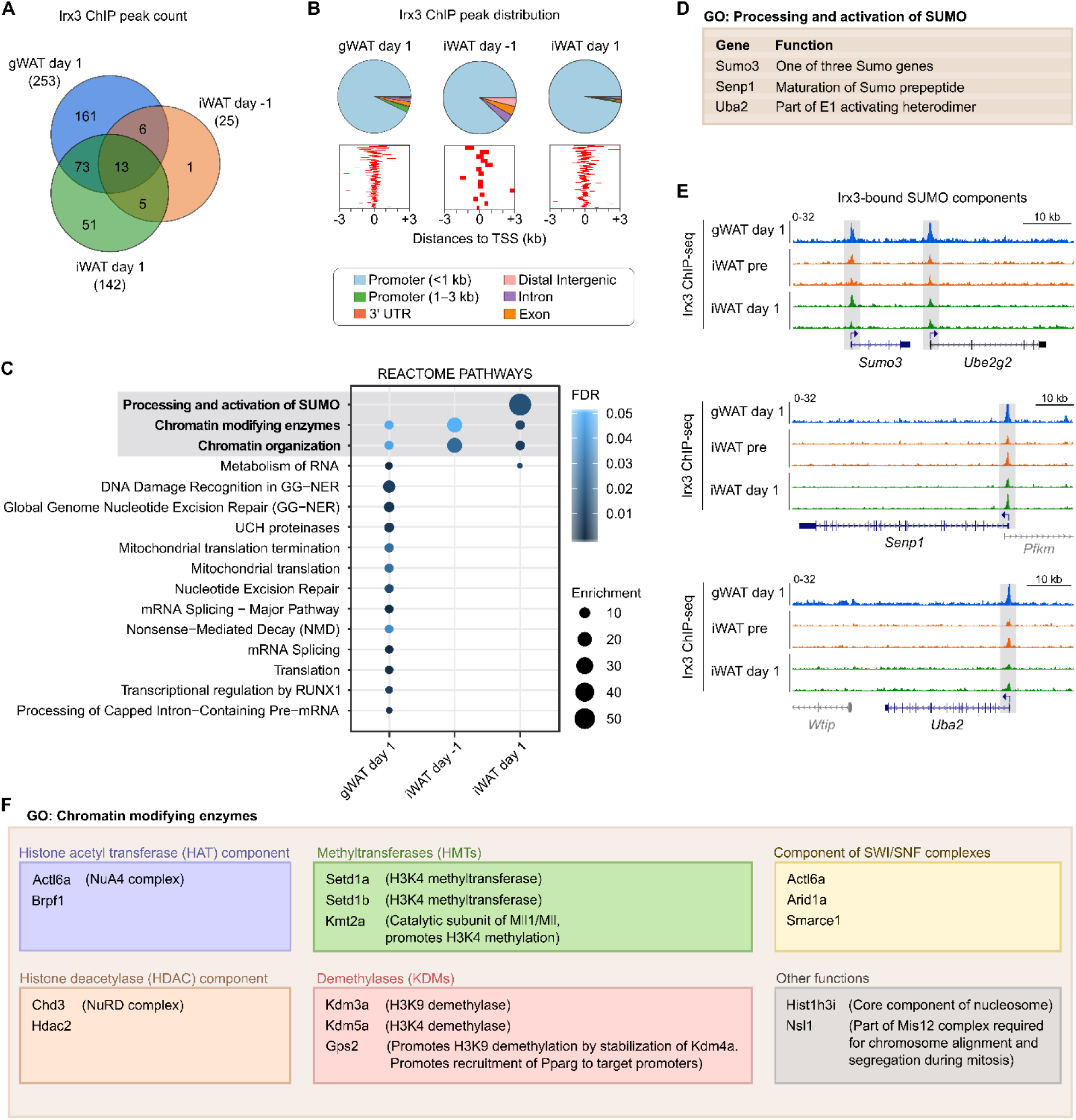
Genome-wide mapping of direct Irx3 target genes in preadipocytes. ChIP-sequencing (ChIP-seq) against Irx3 was performed in preadipocytes from iWAT one day before (day -1) and one day after (day 1) initiation of differentiation, n = 2, as well as from gWAT on day 1, n = 1. A) Number of Irx3 ChIP-seq peaks for each condition shown as a Venn diagram. Only high-confidence (HC) peaks with q-value below 10e-4 and enrichment above 10X were included. B) Distribution of Irx3 ChIP-seq peaks across the genome (top) and relative to nearest TSS (bottom). C) Enriched REACTOME pathway ontology terms for Irx3-bound genes. See also Supplementary file 1 for Panther GOs. D) Genes in the enriched GO category related to Sumoylation. E) Genome browser tracks for each ChIP-seq conditions at the loci of the Sumo-related genes. F) Genes in the enriched GO category related to Chromatin modifying enzymes.

Strikingly, more than 95% of the ChIP-seq peaks were located to proximal promoter regions, with a clear enrichment on transcription start sites (TSSs) (Figure 1B). Thus, virtually all Irx3 binding events could be directly mapped to a target gene (Supplementary file 1). In iWAT, activation of SUMO was the most strongly enriched gene ontology (GO) term, followed by chromatin remodeling (Figure 1C-D and Supplementary file 1). In gWAT, the Irx3 target genes were enriched for myriad GO terms, which also included chromatin modification (Figure 1C and Supplementary file 1). Nonetheless, a more detailed examination of the Irx3 ChIP-seq peaks revealed clear binding of Irx3 to the sumo-related promoters in gWAT as well (Figure 1E). Thus, genes related to processing of SUMO and chromatin modifications were the most consistently enriched Irx3 target genes across both tissues and timepoints. Of note, the promoters of genes encoding both histone acetyl transferases (HATs), deacetylases (HDACs), methyltransferases (HMTs) and demethylases (KDMs), as well as SWI/SNF components were bound by Irx3 (Figure 1F), suggesting a broad, master-regulatory role of Irx3 in epigenetic regulation of preadipocytes.

### Irx3 binds to increasingly open chromatin

We next sought to investigate the chromatin accessibility in Irx3 binding sites during preadipocyte differentiation. To this end, we performed global ATAC-sequencing at seven timepoints during differentiation of preadipocytes from gWAT (Supplementary Figure S1A-D and Supplementary File 2). Integration of ATAC-seq data with the Irx3 ChIP-seq data revealed that 97% and 80% of Irx3 binding events on day 1 occur in regions with >2-fold and >10-fold ATAC enrichment, respectively. Furthermore, these loci remained open throughout differentiation (Supplementary Figure S1E-F), with 40% showing a strong increase in openness in the mature stage, and these were related to adipogenesis and energy homeostasis (Supplementary Figure S1D,G-H). These data suggest that Irx3 plays an important role in the adipogenic program.

Since IRX3 was shown to be important for regulation of white-versus-beige adipocyte identities [12], we further investigated whether the Irx3-bound loci were also accessible in beige adipocytes. We therefore performed ATAC-seq on days 0, 1 and 7 of differentiation in the ME3 beige cell line (Supplementary figure S2). While the total number of ATAC-seq peaks was much lower in the beige compared to white cells, most of the Irx3-bound loci identified in white cells were also open in the beige cells. Moreover, many of these loci also displayed increased openness in the mature cells and were related to stem cell differentiation, histone methylation and transcription. Overall, the chromatin openness of Irx3-bound promoters was highly similar between the white and beige cells, suggesting Irx3 likely controls similar fundamental processes in both cell types independently of its effect on thermogenesis.

### Differentially expressed Irx3 target genes relate to histone modifications and chromatin remodeling

Having identified direct Irx3 target genes by ChIP-seq in WAT, and found similar patterns of chromatin accessibility at the promoters of these genes in both WAT and ME3 cells, we next sought to confirm that these genes are affected by Irx3 depletion. We therefore compared the list of Irx3-bound genes with our previously published list [14] of differentially expressed genes between control and *Irx3*-KO on days 1 and 7 of adipogenic differentiation in ME3 cells. As expected, we found most of the Irx3-bound genes to be sensitive to Irx3 ablation (Figure 2A and supplementary Figure 2). Intriguingly, the direct Irx3-target genes only constituted about 2% of the Irx3-responsive genes (Figure 2A), suggesting that the direct Irx3 target genes act upstream of vast gene networks. In line with this hypothesis, we previously demonstrated that *Irx3-*KO results in loss of adipogenic identity, and this effect was prominent already on day 1 of differentiation [14]. Moreover, in the present work, we found that more than half of the direct Irx3 target genes that were differentially expressed following *Irx3-*KO were affected already on day 1 of differentiation (Figure 2B). These data support an important and early role of Irx3 in determining adipogenic identity.

**Figure 2:**
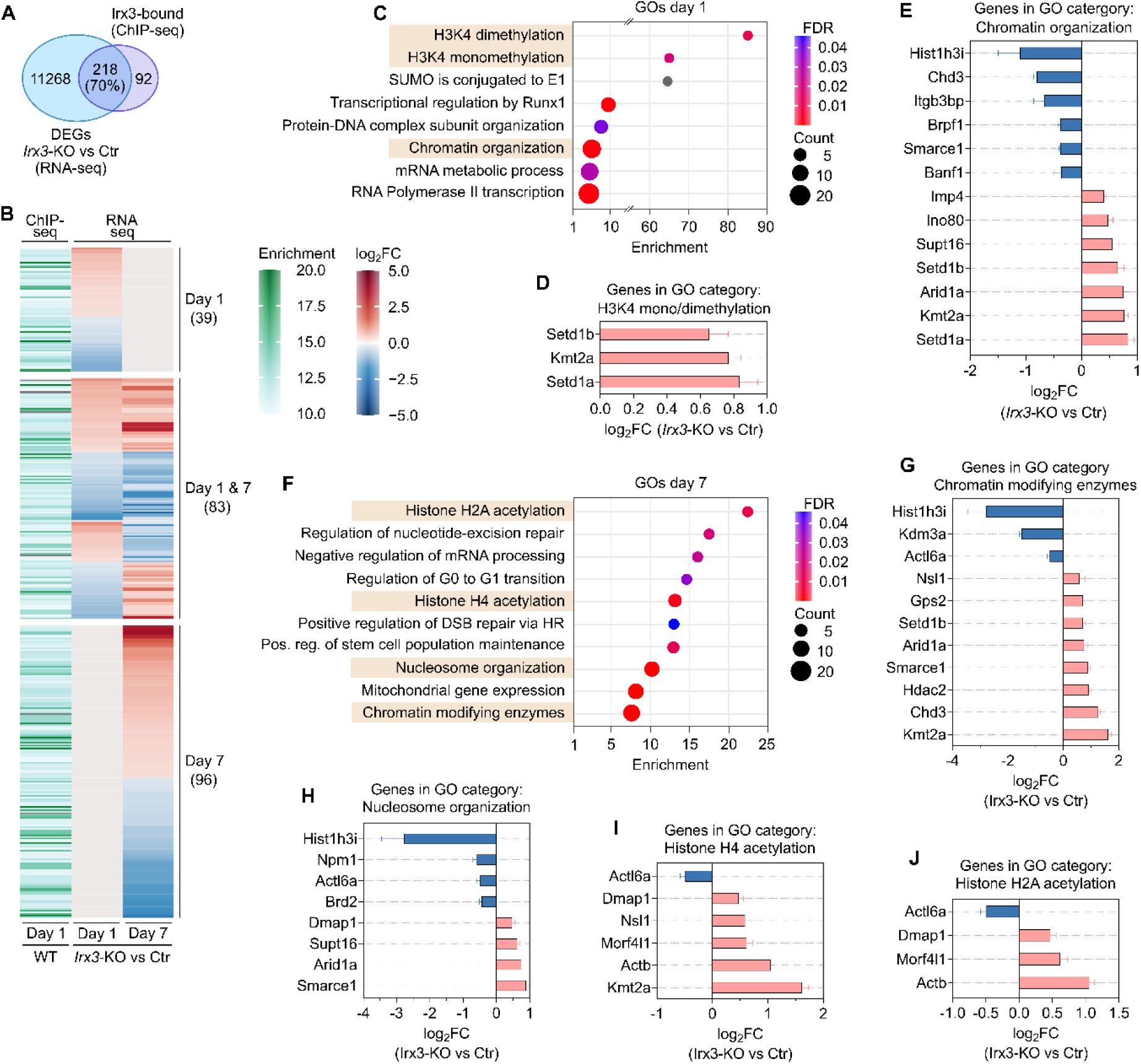
Differentially expressed Irx3 target genes relate to histone modification and chromatin remodeling. A) Venn diagram of Irx3-bound genes and differentially expressed genes in response to *Irx3-*KO. The size of each circle is not drawn according to scale. B) Heatmap displaying Irx3 ChIP-seq peak enrichment and the direction and magnitude of each corresponding Irx3-bound gene in response to Irx3-KO on days 1 and 7 of adipocyte differentiation. C) Enriched Panther and Reactome ontology terms for the 122 Irx3 target genes with differential expression on day 1 of differentiation in response to *Irx3-*KO. D) Genes constituting the GO categories “H3K4 mono/demethylation” on day 1 of differentiation. Log_2_-fold changes for each gene shown. E) Genes constituting the GO category “Chromosome organization” on day 1 of differentiation. Log_2_-fold changes for each gene shown. F) Enriched Panther and Reactome ontology terms for the 179 Irx3 target genes with differential expression on day 7 of differentiation in response to *Irx3-*KO. G) Genes constituting the GO categories “Chromatin modifying enzymes” on day 7 of differentiation. H) Genes constituting the GO categories “Nucleosome organization” on day 7 of differentiation. I) Genes constituting the GO categories “Histone H4 acetylation” on day 7 of differentiation. J) Genes constituting the GO categories “Histone H2A acetylation” on day 7 of differentiation. Log_2_-fold changes for each gene shown. Downregulated genes, blue bars; upregulated genes, red bars. See Supplementary Figure S3 for fold change of genes in the remaining categories.

The top enriched GO categories for day 1 direct target genes were related to histone methylation and chromatin organization, in addition to several other processes (Figure 2C, and supplementary file 3). While the genes encoding histone methyltransferases were consistently upregulated (Figure 2D), the genes involved in chromatin organization were both up- and downregulated (Figure 2E) in *Irx3-*KO cells on day 1. Interestingly, direct Irx3 target genes that were differentially expressed on day 7 were also enriched for histone modifications and chromatin modification, as well as DNA repair and mitochondrial gene expression (Figure 2F-J, supplementary figure S3, and supplementary file 3). Taken together, these data indicate that Irx3 controls the expression of multiple epigenetic factors during both early and late stages of adipogenic differentiation.

### Irx3 represses sumoylation

Having found a broad effect of Irx3 on the expression of epigenetic regulators, we next turned our attention to SUMO, since sumoylation affects the function of epigenetic regulators, and has been shown to be critical for epigenetic regulation of cell identities. While genes involved in sumoylation were enriched in both the ChIP-seq and in the differentially expressed Irx3-bound gene analyses, only the former was significant (Figure 1D and 2C). Moreover, in our initial ChIP-seq analysis, some but not all components of the sumoylation cycle were bound by Irx3. However, because we used strict criteria in the untargeted ChIP-seq analysis, we hypothesized that other sumo components might also be bound, albeit weaker. Manual curation of the ChIP-seq tracks revealed that, at least in gWAT, genes in all parts of the sumoylation cycle were indeed bound to some degree by Irx3 (Figure 3A).

**Figure 3:**
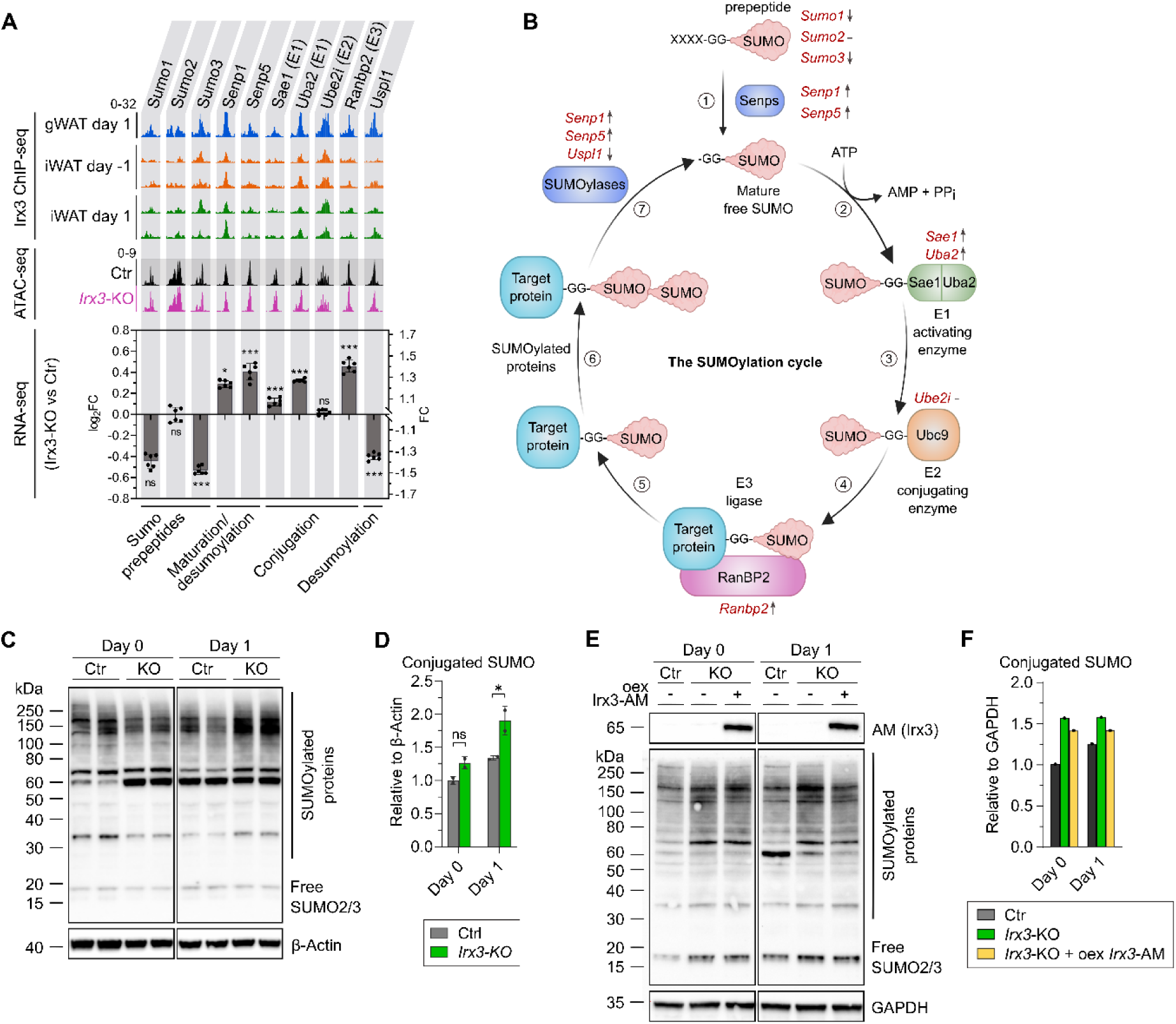
Irx3 represses sumoylation. A) Irx3 ChIP-seq peaks at the promoter (TSS ±1.5 kb) of key genes involved in sumoylation one day before and/or one day after initiation of adipogenic differentiation in preadipocytes isolated from iWAT and gWAT (top). ATAC-seq at the same loci in ME3 control and *Irx3-*KO cells on day 1 of adipogenic differentiation (middle). Differences in expression of the respective genes between ME3 control and *Irx3-*KO cells on day 1. Log_2_-fold changes and fold changes shown (bottom). B) Overview of the sumoylation cycle. The genes shown in (A) are highlighted in red letters with arrows indicating direction of change with *Irx3-*KO. The *Sumo* genes encode prepeptides that can be cleaved and activated by Senp proteases to reveal a GG diglycine motif (1). The GG motif in free, mature SUMO forms a thioester bond with the E1 activating complex, consisting of the Sae1 and Uba2 (also known as Sae2) heterodimer (2). The SUMO group is next transferred to the E2 conjugating enzyme Ubc9 (also known as Ube2i) (3), which in turn transfers the SUMO group to the target protein with the aid of E3 ligases (4-5). Multiple SUMO subunits can be attached together to form a poly-SUMO tail (6). SUMO groups can be released from target proteins by Senp or other SUMOylase family members (7). C) Western blot showing global SUMOylome pattern on days 0 and 1 of adipogenic differentiation in ME3 control and *Irx3-*KO cells. D) Volumetric quantification of bands in (C). Levels of conjugated SUMO relative to β-Actin shown. \**p* < 0.05; ns, not significant, two-way ANOVA with Holm-Sidak correction for multiple testing. E) Western blot showing global SUMOylome pattern on days 0 and 1 of adipogenic differentiation in ME3 control and *Irx3-*KO cells transfected with either empty plasmid or AM-tagged Irx3. F) Volumetric quantification of bands in (E). Relative levels of conjugated SUMOylome relative to GAPDH shown.

Moreover, most of these genes were differentially expressed in response to *Irx3-*KO, and this effect was largely independent of chromatin openness (Figure 3A). Specifically, both *Senp1* and *Senp5* required for maturation of Sumo, as well as the *Sae1* and *Uba2* components of the E1 activating enzyme and Ranbp2, an E3 ligase, were all upregulated (Figure 3A-B). Conversely, the desumoylase *Uspl1* was downregulated in *Irx3-*KO cells. These data suggest a repressive role of Irx3 on sumoylation and thereby increased conjugation of SUMO in the *Irx3-*KO cells. However, because *Sumo3* itself also was downregulated, the overall effect of Irx3 on sumoylation was not entirely clear. Thus, to functionally test whether Irx3 could suppress or promote sumoylation, we differentiated control and *Irx3-*KO ME3 cells for 0, 1 and 7 days and lysed them under strict denaturing conditions in the presence of the desumoylase inhibitor N-ethylmaleimide (NEM) to preserve SUMO conjugates.

Western blotting using a Sumo2/3 antibody revealed increased levels of conjugated SUMO in the absence of Irx3 (Figure 3C-D). Furthermore, reintroducing Irx3 in the KO cells reduced conjugated SUMO levels (Figure 3E-F). These data confirm that Irx3 suppresses sumoylation during early differentiation of beige ME3 adipocytes.

### Inhibition of sumoylation restores adipogenesis in *Irx3-*KO cells

We previously demonstrated that loss of Irx3 in beige cells abolishes adipogenesis [14], and we show here that Irx3 ablation also increases sumoylation in the same cell type. Because sumoylation has been reported to repress adipogenesis [30,31], we hypothesized that elevated sumoylation may mediate the repressive effect of Irx3 ablation on adipogenesis. If so, pharmacological inhibition of sumoylation should restore adipogenesis in the *Irx3-*KO cells. To test this, we treated control and *Irx3-*KO cells with the sumoylation inhibitor ML-792 from days -2 to 9 of differentiation and assessed the effect on differentiation (Figure 4). Strikingly, ML-792 treatment had a profound impact on the differentiation capacity of the *Irx3-*KO cells (Figure 4A-B and supplementary figure S5A), and was able to completely restore the total lipid levels (Figure 4C). This effect was attributable to both a partial restoration of the number of differentiating cells, as well as increased lipid droplet size in each cell (Figure 4C). In accordance with this, the expression of both the adipogenic master regulators Pparγ and C/ebβ, as well as other markers of mature adipocytes and lipid metabolism were increased in response to ML-792 (figure 4D and Supplementary figure S5B). Of note, ML-792 also had a mild proadipogenic role in the control cells, but the effect was always more profound in the *Irx3-*KO cells. A notable exception was Ucp1, which was stimulated in the control cells, but not in the *Irx3-*KO cells. ML-792 demonstrated a dose-response effect on lipid accumulation in the *Irx3-*KO cells (Figure 4E and Supplementary figure S5C), with minimal toxicity at 0.5 µM (Supplementary figure S5D).

**Figure 4:**
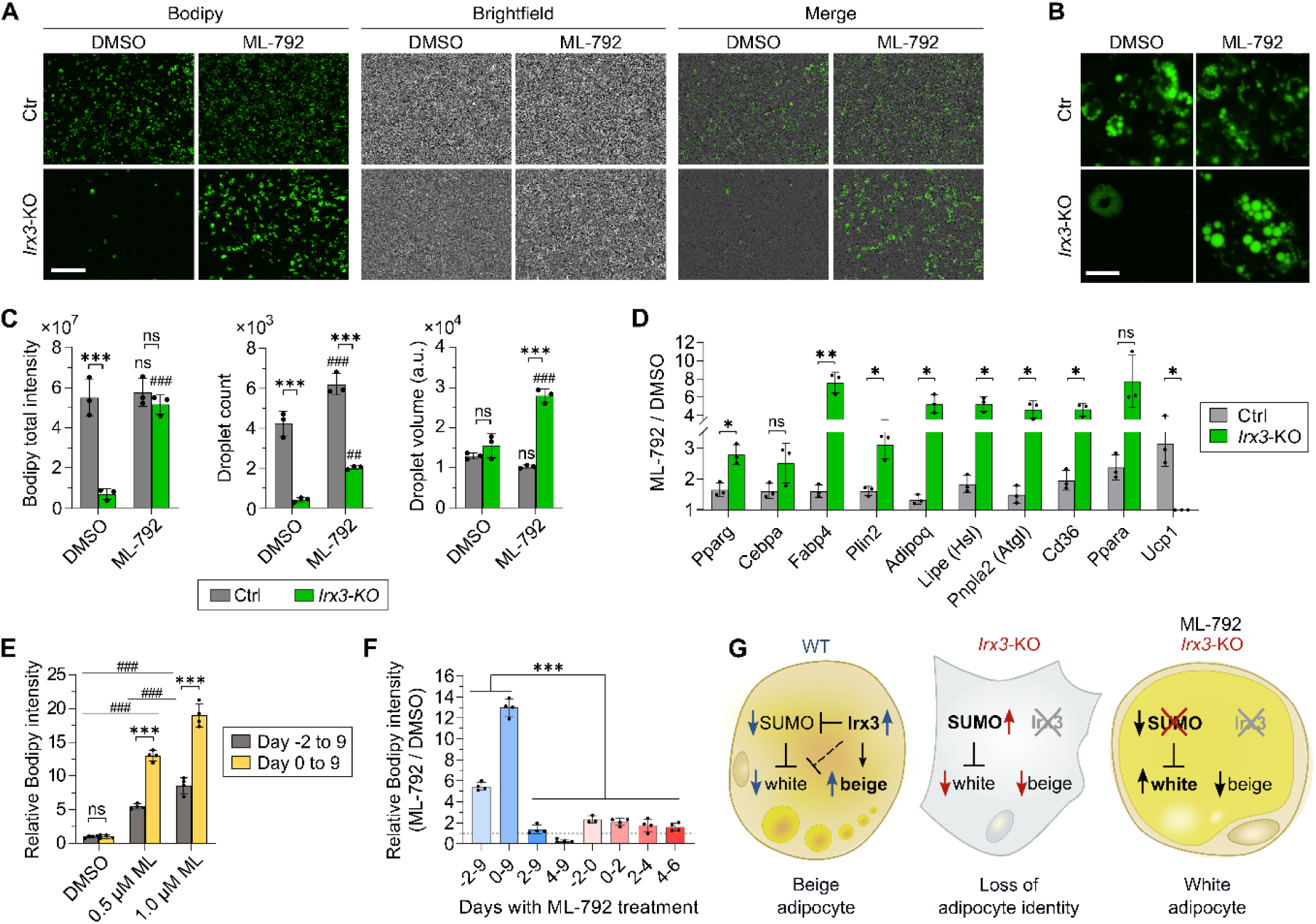
Inhibition of sumoylation restores adipogenesis in *Irx3-*KO cells. A) ME3 control and *Irx3-*KO cells were treated with either vehicle or 0.5 µM of the sumoylation inhibitor ML-792 from days-2 to 9 of adipogenic differentiation. On day 9, the cells were treated with the Bodipy fluorescent lipid stain for 1 hour and imaged with fluorescence and brightfield microscopy. Representative images from 9 images/well in biological triplicates, from one of four independent experiments. Scale bar = 400 µm. See Supplementary figure S5A for all images from this experiment. B) Magnification of the images shown in (A). Scale bar = 50 µm. C) Quantification of Bodipy total integrated intensity, number of lipid droplet and average lipid volumes. Data from all 9 images/well were merged to represent each well. Three replicate wells from one of four independent experiments are shown. Data normalized to relative cell count. ***p < 0.001, control vs *Irx3-*KO cells; ^##^p <0.01, ^###^p < 0.001, ML-792 vs DMSO; ns, not significant, two-way ANOVA with Holm-Sidak correction for multiple testing. D) Gene expression in ME3 control and *Irx3-*KO cells in response to 0.5 µM ML-792, same cells as (A-C). Ratios between ML-792 and DMSO for control and *Irx3-*KO cells shown for simplicity. Rps13 was used as reference gene. See Supplementary figure S5B for full gene expression data. *p < 0.05, **p < 0.01, multiple Student’s t-tests with Holm-Sidak correction for multiple testing. E) Dose-response effect of ML-792 on lipid accumulation in *Irx3-*KO cells when ML-792 was added on either day -2 or 0 and kept throughout differentiation until day 9. Data from 4 images/well merged to represent each well. Quantification of four replicate wells from one of three independent experiments shown. See supplementary figure S5C for images. *p < 0.05, ***p <0.001, ^###^p < 0.001; ns, not significant, two-way ANOVA with Holm-Sidak correction for multiple testing. F) The effect of keeping 0.5 µM ML-792 at different stages of differentiation (blue bars), or only during various 2-day intervals (red bars). Data normalized to relative cell count and shown as ML-792/DMSO ratio. ***p < 0.001, one-way ANOVA with Holm-Sidak correction for multiple testing. G) Schematic with proposed mechanism for Irx3 and Sumo-mediated regulation of adipogenesis.

At both 0.5 µM and 1.0 µM of the inhibitor, the pro-adipogenic effect was roughly doubled when administered from day 0 instead of day -2 (Figure 4E), indicating the importance of inhibiting sumoylation *after* the induction of adipogenesis. These results prompted us to test whether the pro-adipogenic effect of ML-792 was acting during early or late stages of differentiation. Strikingly, delaying the administration of the inhibitor to day 2 or later completely abolished the pro-adipogenic effect, and administration from day 4 actually repressed rather than stimulated differentiation (Figure 4F). These data indicate that inhibition of sumoylation plays a particularly important role during days 0-2 of differentiation, i.e., in the window where variants in the *FTO* locus were previously shown to affect IRX3 expression [12]. However, treatment with ML-792 solely on days 0-2 resulted in only a minor stimulatory effect (Figure 4F), suggesting that inhibition of sumoylation during the two first days of differentiation is necessary, but not sufficient to promote adipogenesis. Taken together, these data clearly demonstrate that inhibition of sumoylation during adipogenic stimulation rescues the adipogenic capacity of *Irx3-*KO cells, but only in the direction of white and not beige adipocytes. This suggests that elevated sumoylation mediates some, but not all the effects of Irx3 ablation in the ME3 cells (Figure 4G).

### Inhibition of sumoylation improves Pparγ transcriptional activity and its synergy with Pgc-1α

The dramatic effects of Irx3 and sumoylation on adipogenesis suggest that these factors control key regulators in adipogenesis, such as Pparγ. Surprisingly, although *Pparg* was indeed strongly downregulated during late stages of adipogenic stimulation in *Irx3-*KO cells [14], its expression on day 1 was not reduced but rather slightly upregulated (Figure 5A). However, downstream Pparγ target genes, which themselves are adipogenic activators such as Ebf2, Pgc-1a and Prdm16, were strongly downregulated already on day 1 (Figure 5A). This suggests impaired Pparγ *function* following loss of Irx3. Since sumoylation generally has been found to have a repressive effect on target protein function and Pparγ is a known SUMO target (Figure 5A), we hypothesized that the elevated sumoylation following *Irx3-*KO could repress Pparγ activity. To test this theory, we tested the ability of Pparγ and its co-activator Pgc-1α to activate a luciferase reporter under control of an artificial Ppar-response element (PPRE) in ME3 cells with or without ML-792 (Figure 5B). Indeed, inhibition of sumoylation increased Pparγ transcriptional activity 2-4-fold, and this effect was observed both with and without the synthetic ligand rosiglitazone (rosi), although the overall strongest effect of ML-792 was observed in the presence of both rosi and Pgc-1α (Figure 5B).

**Figure 5:**
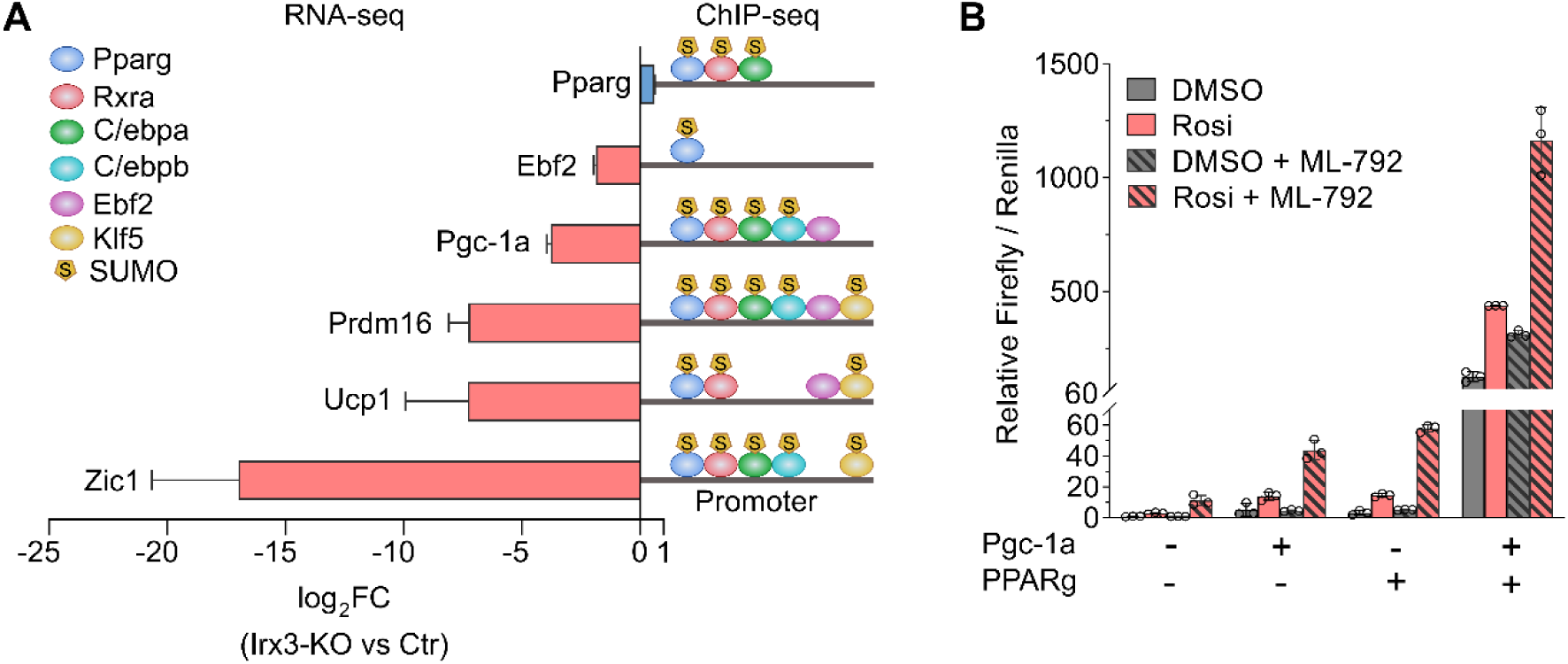
Inhibition of sumoylation improves Pparg transcriptional activity and its synergy with Pgc-1a. A) Differential expression of adipogenic regulators in *Irx3-*KO vs control cells. Experimentally validated binding of upstream transcription factors to the promoters of each adipogenic regulator in WT cells is shown to the right, ChIP-seq data collected from the UCSC genome browser hub UniBind 2021 [32]. Transcription factors experimentally shown to be sumoylated in WT preadipocytes or mature adipocytes are marked with (S), data from [33]. B) Luciferase activity of a reporter construct under control of 3xPPRE sites co-transfected with Pparg and/or Pgc-1a. Firefly luciferase units relative to the control group and normalized to constitutive Renilla luciferase is shown. Data show mean ± SD of n= 3 replicates from one out of two independent experiments.

### Irx3 and SUMO share common target genes involved in adipocyte versus osteoblast development

To better understand which Irx3-dependent processes are mediated via sumoylation, we next compared the changes in global gene expression in response to *Irx3-*KO and ML-792 (Figure 6). We first took advantage of our recently published SLAM-seq dataset on the effect of ML-792 *on newly transcribed* genes on day 1 of differentiation in 3T3-L1 cells [33]. Of note, ME3 and 3T3-L1 cells exhibit similar gene expression profiles, particularly during early stages of differentiation. While only 471 newly transcribed genes responded to ML-792 using a fold change cut-off of 1.2, more than half of these genes were also differentially expressed in response to *Irx3-*KO (Figure 6A). Interestingly, genes upregulated in both datasets were functionally enriched for positive regulation of histone modifications, while common downregulated genes were enriched for negative regulation of osteoblast differentiation (Figure 6B-C) in addition to a range of morphogenic processes (Supplementary Figure S6 and Supplementary file 4).

**Figure 6:**
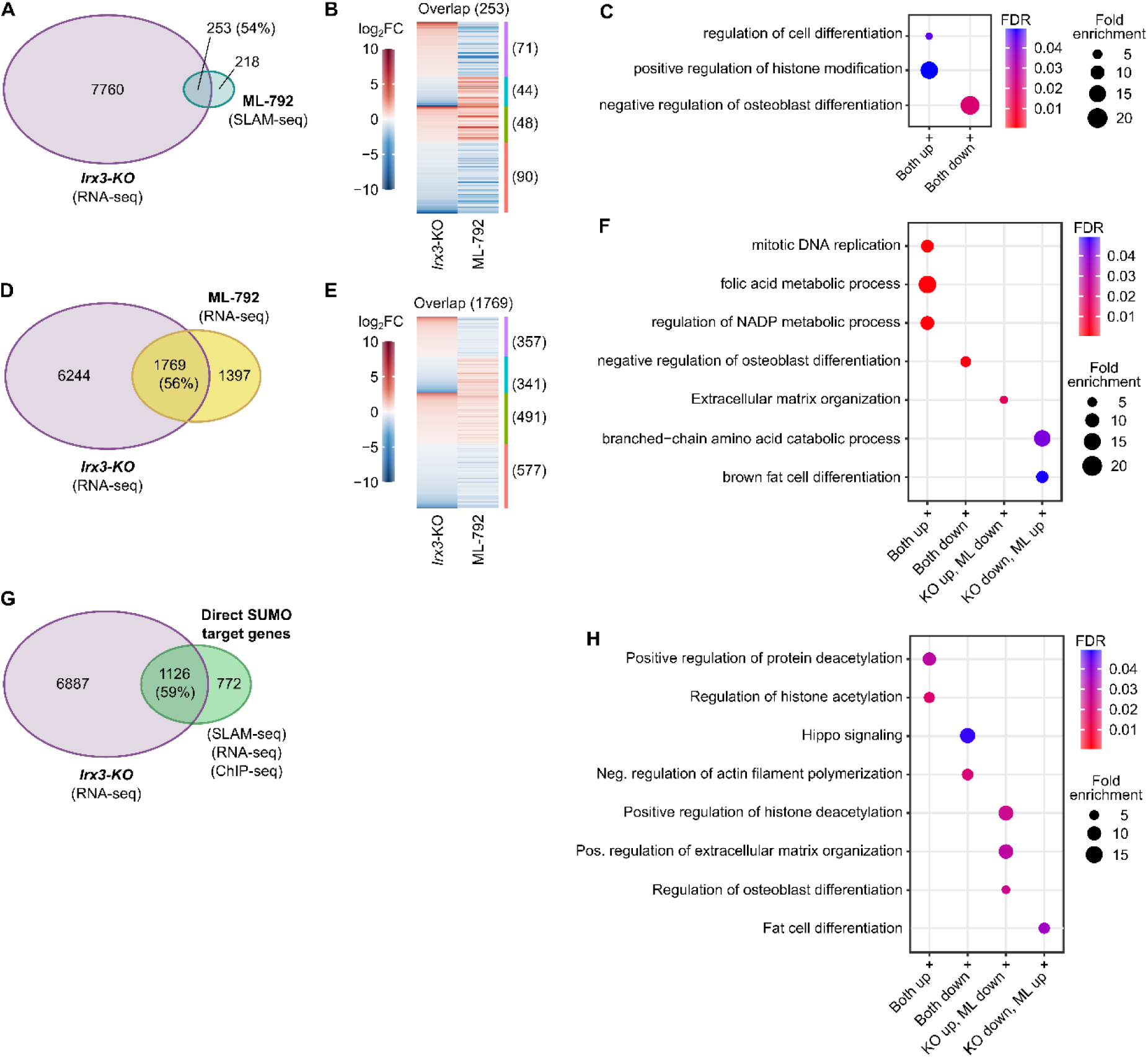
Irx3 and SUMO share common target genes involved in adipocyte versus osteoblast development. Global mRNA-sequencing was performed to map differentially expressed genes (DEGs) between ME3 control and *Irx3-*KO cells on day 1 of differentiation. This list was compared with either differences in *newly transcribed* genes (DNTs) as measured by SLAM-seq (A-C), or DEGs as measured by mRNA-seq (D-F) in response to ML-792 in 3T3-L1 cells. A) Venn diagram showing overlap between *Irx3-*KO mRNA-seq DEGs and ML-792 SLAM-seq DNTs on day 1 of differentiation. B) Heatmap showing direction of change among overlapping genes in (A). C) Selection of top enriched/most significant GO categories among the overlapping genes in (A-B). See Supplementary Figure S6 and Supplementary file 4 for complete lists. D) Venn diagram showing overlap between mRNA-seq DEGs in response to *Irx3-*KO or ML-792 on day 1 of differentiation. E) Heatmap showing direction of change among overlapping genes in (D). F) Selection of top enriched/most significant GO categories among the overlapping genes in (D-E). See Supplementary Figure S7 and Supplementary file 5 for complete lists. G) A pooled list of genes responding to ML-792 identified by SLAM-seq and mRNA-seq was overlapped with a SUMO-ChIP-seq dataset to make a list of direct SUMO target genes (Supplementary file 6), which in turn was compared with the list of Irx3-responsive genes. H) Selection of top enriched/most significant GO categories among the overlapping genes in (G). See Supplementary file 6 for complete lists.

While the SLAM-seq provides an intriguing snapshot image of genes responding to ML-792 on day 1 of differentiation, it is not designed to take into account accumulated changes in mRNA levels over time. However, we previously also measured the effect of ML-792 on 3T3-L1 cells using RNa-seq [33]. Therefore, to directly compare differentially expressed genes in response to ablation of Irx3 or inhibition of sumoylation, we next compared RNA-seq data from both experiments (Figure 6D-F). As expected, the number of overlapping genes was larger, but the proportion of ML-792 sensitive genes that were also responsive to *Irx3-*KO was similar between RNA-seq and SLAM-seq (Figure 6D). We found sumoylation and Irx3 to regulate a shared set of genes involved in (brown) fat cell differentiation in opposite directions (Figure 6F), in agreement with our data showing a repressive effect of Irx3 on sumoylation during adipogenesis and inverse effects of Irx3 and sumoylation on adipogenesis.

We also compared the response in global gene expression to lack of Irx3 or sumoylation inhibition on day 7 of differentiation (Supplementary figure S8). In the mature adipocytes, 70% of sumoylation-sensitive genes were also affected by *Irx3-*KO, and “sumoylation of transcription factors” was among the top enriched/most significant GO terms for genes repressed in the absence of Sumo, but activated in the absence of Irx3. Thus, these data confirmed the repressive effect of Irx3 on sumoylation.

While we have identified pathways affected by ablation of both Irx3 and sumoylation, we next sought to determine whether we could identify the *direct* SUMO target genes and compare them with Irx3-responsive genes. To this end we first pooled the ML-792 responsive genes identified from the SLAM-seq and RNA-seq datasets, using significantly changed genes, but with no fold change filter (Supplementary file 6). We then compared this extended list of genes with our recently published Sumo2/3 ChIP-seq dataset [33] to identify genes that are bound by sumoylated transcription factors and that change expression in response to ML-792 (Supplementary file 6). These direct SUMO target genes were subsequently compared with the *Irx3-*KO dataset (Figure 6G). Nearly 2/3 of the direct SUMO target genes overlapped with the *Irx3-*KO responsive genes, and oppositely regulated genes were enriched in GO terms related to differentiation of both adipocytes and osteoblasts, in addition to regulation of histone deacetylation (Figure 6H and Supplementary file 6). Taken together, our data show that a majority of direct SUMO target genes are regulated by Irx3, strengthening our finding of Irx3 as an upstream regulator of Sumo. Moreover, the shared target genes are involved in a range of biological processes, including histone modifications and regulation of adipogenesis and osteogenesis.

### *Irx3-*KO promotes osteogenesis

Our gene expression data suggest that both Irx3 and sumoylation may be involved in adipocyte versus osteoblast determination. Specifically, *Irx3-*KO cells appeared to have lost their resistance towards osteoblast differentiation. We hypothesized that this effect could reflect an underlying alteration of the epigenetic landscape, so we performed ATAC-seq to measure the differences in open chromatin during adipogenic differentiation of the control and *Irx3-*KO cells. More than 35,000 genomic loci displayed differential openness in response to Irx3 ablation on day 0, and this number was increased to 93,000 and 81,000 on days 1 and 7, respectively (Supplementary file 7), demonstrating a profound effect of Irx3 on the global chromatin landscape. Since we did not have RNA-seq data on day 0, we were particularly interested in the ATAC-seq data for this timepoint as it would reflect the developmental potential of the *Irx3-*KO cells before being subjected to adipogenic stimulation. Indeed, GO analyses of ATAC-seq peaks in more open promoters on day 0 revealed significant enrichment of several processes related to Wnt-signaling, osteoblast differentiation and bone generation (Figure 7A-B, Supplementary figure S9 and supplementary file 8), supporting an epigenetic and developmental shift in the *Irx3-*KO cells. Moreover, similar GO terms were found for more open chromatin on days 1 and 7 (Supplementary figure S9 and supplementary file 8), suggesting that the pro-osteogenic changes present on day 0 remained and resisted adipogenic stimulation. Finally, we observed strong concordance between the RNA-seq and ATAC-seq data (Supplementary figure S10 and supplementary file 9). Overall, these data indicate a more permissive epigenetic landscape towards osteogenesis in the *Irx3-*KO cells.

**Figure 7:**
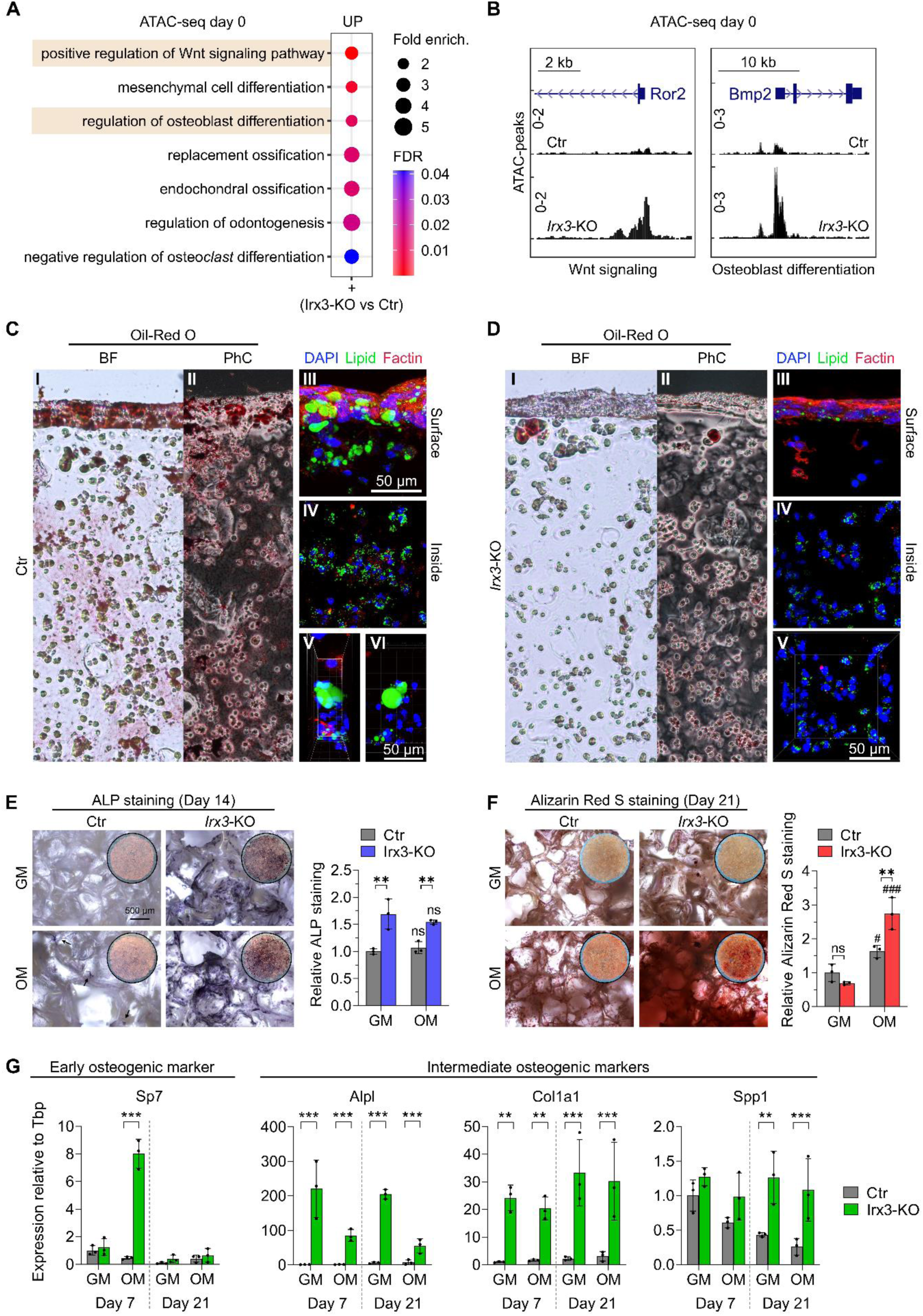
*Irx3-*KO promotes osteogenesis. A) Open chromatin profiling was performed by ATAC-seq in control and *Irx3-*KO cells on day 0 of differentiation. Selected top GO for more open peaks in promoters of *Irx3-*KO cells shown. See also supplementary figure S9. B) UCSC genome browser tracks showing examples ATAC-seq peaks from control and *Irx3-*KO cells at the promoters of genes from two osteoblast-related GO categories in (A). C) ME3 control cells subjected to adipogenic differentiation in 3D culture. I, brightfield view of Oil-red-O lipid-stained cells. II, phase contrast view of Oil-red-O staining. III, fluorescent view of the hydrogel surface with DAPI nuclear stain (blue), Bodipy lipid stain (green) and F-actin cytoskeletal stain (red). IV, fluorescent view of the inside of the hydrogel. V and VI, zoomed in fluorescent view. D) *Irx3-*KO control cells subjected to adipogenic differentiation in 3D culture. I-V, same conditions as in C. E) ME3 control and *Irx3-*KO cells treated with growth medium (GM) or osteogenic medium (OM), followed by Alkaline Phosphatase (ALP) staining. Brightfield view (left panel) and quantification (right panel) of ALP staining. F) ME3 control and *Irx3-*KO cells treated with growth medium (GM) or osteogenic medium (OM), followed by Alizarin Red S staining (ARS). Brightfield view (left panel) and quantification (right panel) of the ARS staining. G) Expression of early, intermediate and late osteogenic markers in in control and *Irx3-*KO cells in response to differentiation in growth medium or osteogenic medium. Two-way Anova with Holm-Sidak correction for multiple testing. # *p_adj_* < 0.05, ### *p_adj_* < 0.001, osteogenic vs growth medium. \**p*_adj_ < 0.05, \*\**p*_adj_ < 0.01, \*\*\**p*_adj_ < 0.001 control vs *Irx3-*KO. NS, not significant.

To experimentally test whether the epigenetic propensity towards osteogenesis in the *Irx3-*KO cells could permit these cells to undergo osteogenesis under the proper conditions, we prepared 3D cultures of the ME3 cells and subjected them to either adipogenic (Figure 7C-D) or osteogenic (Figure 7E-F) stimulation. As expected, the ME3 control cells readily differentiated into mature adipocytes with larger lipid droplets formed following adipogenic stimulation (Figure 7C) while the *Irx3-*KO cells did not, as assessed by cell morphology and Oil-red-O and Bodipy lipid staining (Figure 7D).

Conversely, while the control cells showed no or minimal response to osteogenic stimulation, the *Irx3-*KO cells demonstrated significantly higher basal or induced levels of osteogenic Alkaline Phosphatase (ALP) (Figure 7E) and matrix mineralization confirmed by Alizarine Red S (Figure 7F) staining. Gene expression data revealed a potent increase in a key transcription factor for osteogenic differentiation, *Sp7,* on day 7 of osteogenic stimulation in *Irx3-*KO compared to control cells (Figure 7G). In contrast, the *Runx2*, *Sost* and *Bglap* markers were either not significantly affected or not expressed in the KO cells (data not shown). However, the intermediate osteogenic marker *Spp1* was elevated in the *Irx3-*KO cells on day 21 in both growth medium and the osteogenic medium, while the intermediate osteogenic markers *Alpl* and *Col1a1* were strongly upregulated in the *Irx3-*KO cells at both timepoints and in both media (Figure 7G). Taken together, the *Irx3-*KO cells demonstrate an altered epigenetic landscape that promotes osteogenic over adipogenic differentiation.

### Inhibition of sumoylation represses *Irx3-*KO-dependent osteogenesis

To assess whether increased sumoylation levels could mediate the pro-osteogenic effect of Irx3 ablation, we treated 3D cultures of control and *Irx3-*KO cells grown in osteogenic medium with either vehicle or the ML-792 inhibitor of sumoylation (Figure 8). The *Irx3-*KO cells treated with vehicle showed elevated mineralization and expression of osteogenic markers compared to control cells, while inhibition of sumoylation significantly decreased both mineralization and osteogenic gene expression in the KO cells. These data suggest that sumoylation at least partially mediates the pro-osteogenic effect of Irx3 ablation.

**Figure 8:**
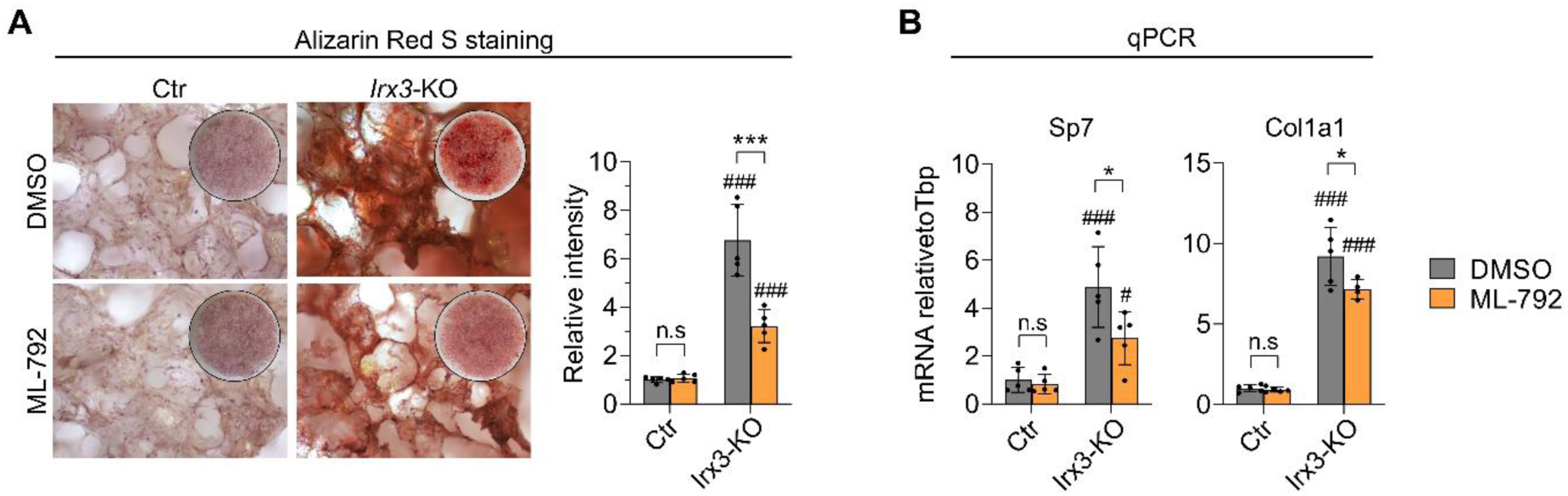
Inhibition of sumoylation represses *Irx3-*KO-dependent osteogenesis. A) ME3 control and *Irx3-*KO cells were subjected to osteogenic differentiation in 3D culture and treated with either vehicle or 0.5 µM ML-792. On day 21, the cells were stained with alizarin Red S. Brightfield view (left panel) and quantification (right panel) shown. Data represent n=5 replicate wells from a single experiment. B) Cells were treated as in (A), except RNA was harvested on day 7 and expression of early and intermediate markers of osteogenesis was measured by qPCR. Data shown relative to the reference gene *Tbp* and the control group. Data represent n=5 replicate wells from a single experiment. Two-way Anova with Holm-Sidak correction for multiple testing. # *p_adj_* < 0.05, ### *p_adj_* < 0.001, control vs *Irx3-*KO. \**p*_adj_ < 0.05, \*\*\**p*_adj_ < 0.001 DMSO vs ML-792. Nns, not significant.

## Discussion

The present study has identified hundreds of novel direct target genes of the homeobox transcription factor Irx3 in preadipocytes, providing mechanistic insight into how IRX3 mediates the association between risk variants in the *FTO* locus and increased risk of obesity [12][34]. We previously found a transient, risk genotype-dependent increase in IRX3 during early adipogenesis to promote white over beige adipocyte identity of the mature adipocytes [12,35]. These data suggest an epigenetic effect of Irx3 and despite its lack of chromatin or histone-modifying domains we here found that chromatin remodeling and histone modifying enzymes were the top enriched biological pathways among the identified direct Irx3 target genes. Moreover, these genes were differentially expressed following *Irx3-* KO, showing that Irx3 directly controls their expression. Strikingly, Irx3 was found to transcriptionally suppress the sumoylation machinery and reduce global conjugated levels of SUMO, a key PTM involved in modulating the function of chromatin remodelers and histone modifiers [24–27]. Taken together, these data indicate that Irx3 controls both the expression and function of epigenetic regulators, and implicate Irx3 as a novel upstream regulator of sumoylation In agreement with a role of Irx3 in transcriptionally controlling epigenetic modifiers, we identified profound changes in the open chromatin landscape in *Irx3-*KO compared to control cells, with more open chromatin in promoters of osteogenic genes and less open chromatin in adipogenic promoters. Consequently, the *Irx3-*KO cells failed to undergo adipogenesis both in 2D and 3D culture. Conversely, we found the *Irx3-*KO cells to respond to osteogenic stimulation, while the control cells did not. Finally, pharmacological inhibition of sumoylation partially reversed the *Irx3-*KO dependent switch from adipogenesis to osteogenesis. SUMO is well known to control cell fates, and have been shown to be critical in distinguishing somatic, differentiated cells from induced pluripotent stem cells, embryonic stem cells and 2C-like cells [28,29]. In the present study, we found Irx3 to be an upstream transcriptional regulator of epigenetic remodelers, and able to control cell fate at least in part via SUMO.

Identification of genes influencing obesity and other complex diseases has long been hampered by the lack of mechanistic information from GWAS data [3,9,36]. While a major leap was achieved by us and others a decade ago by inferring a causal variant in the obesity-associated *FTO* locus, identifying IRX3 and IRX5 as the target genes and adipocyte precursor cells as the effector cell type [12], the question still remained how these genes mechanistically mediate the effect of the association [3]. We did find a transient, genotype-dependent increase in IRX3 and IRX5 expression on days 0-2 of adipogenic differentiation in adipocyte precursor cells, which subsequently inhibited beiging and promoted a white phenotype in mature adipocytes [12]. However, the exact nature of this temporal effect was not investigated. While more recent work by Vámos et al. confirmed reduced thermogenic capacity in human adipose-derived precursor cells from patients homozygous for the *FTO* locus risk variants [35], the authors did not report the expression levels of IRX3. Moreover, Zou et al. reported a conflicting effect of IRX3 on adipocyte beiging and found IRX3 to promote rather than inhibit beiging in mice and humans [37]. Thus, there has been a need for more mechanistic investigations into the role of IRX3 in adipocytes [38]. Zou et al. found reduced expression of thermogenic markers like UCP1 in beige adipocytes following lentiviral-mediated knockdown of IRX3. Therefore, if IRX3 primarily acts through uncoupling, this finding should lead to *increased* lipid accumulation in the *IRX3-*kd cells. Instead, they found *reduced* lipid accumulation in the *IRX3-*kd cells [37]. We reasoned the repressive effect on beige gene expression could be confounded by a suppressive effect on adipogenesis *per se*. Indeed, using CRISPR/Cas9 to completely knock down Irx3 in beige cells, we previously demonstrated a near complete inhibition of adipogenesis [14]. The current study broadened this finding, showing that Irx3 is vital for maintaining an epigenetic landscape and transcriptional program tuned towards adipogenesis, while simultaneously inhibiting chondrogenesis and osteogenesis. This positions Irx3 as a master regulator of mesenchymal fate, and provides an important new step in elucidating the biological mechanisms underlying the *FTO* locus’ association with obesity. Interestingly, a recent study showed that in hypertrophic chondrocytes of mice, Irx3 and Irx5 promoted transdifferentiation towards osteogenesis and inhibited adipogenesis [39]. On the other hand, another study found Irx3 to promote chondrogenesis from mesenchymal stem cells in the presence of Bmp2 [40]. While the direction of regulation appears context-specific, these studies underscore the role of Irx3 as a key regulator of mesenchymal fat decision.

Although the direct Irx3 target genes may vary between different cell types and tissues, many of the genes identified in preadipocytes are nevertheless involved in biological pathways present in multiple cell types. For example, our data show that Irx3 controls genes in the Wnt/β-catenin and Hippo signaling pathways in preadipocytes, and these pathways have been shown by others to also be regulated by Irx3 and/or Irx5 in kidney development [41]. Thus our data may help understand how IRX3 mediates its effects in several other conditions in which IRX3 has been implicated, including type 2 diabetes [10], metabolic inflammation [42], fertility [43], neurogenesis [44,45], glioblastoma [46], acute myeloid leukemia [47–50], microvascularization [51] and heart development and function [52– 57].

The master-regulatory role of Irx3 may not be surprising given its classification as a homeobox factor [15,58,59], a highly diverse super-class of transcription factors with critical roles during development, patterning formation and embryonic stem cell function [15,16]. Interestingly, SUMO has been reported to regulate the function of several homeobox factors, including *Nkx2.5* via its direct sumoylation [60] and *Egl-5* and *Mab-5* via sumoylation of the Sop-2 Polycomb repressor protein [61]. However, the present report is the first to show a homeobox factor acting *upstream* of SUMO, and our discovery fills an important general knowledge gap as no upstream SUMO regulator has been identified previously (at least to our knowledge). This finding should stimulate further studies of the upstream mechanisms controlling SUMO and warrants investigations into the role of Irx3 in a broad range of known SUMO-controlled processes, including stemness and cell identity, chromatin and epigenetic remodeling, stress response, cell division, senescence, protein stability and more, as reviewed in [26]. Moreover, potential implications for diseases where SUMO is involved, such as acute promyelocytic leukemia (APL) and neurodegenerative diseases (Alzheimer’s, Parkinson’s and Huntington’s disease) [26] remains to be examined.

In the present study we found sumoylation to have a strong inhibitory role on adipogenesis, which could be reversed by pharmacological inhibition of sumoylation. This finding contrasts some other reports suggesting that SUMO promotes adiposity. For instance, Mikkonen et al. found *Sumo1-*KO mice fed a high-fat diet to weigh less and possess smaller and fewer adipocytes compared to WT mice [62]. Similarly, mice lacking the SUMO conjugating enzyme Ubc9 specifically in WAT were severely lipodystrophic [63] and *Ubc9*-KO in 3T3-L1 cells inhibits adipogenesis [64]. Interestingly, mouse embryonic fibroblasts (MEFs) with *Sumo1* deficiency demonstrated reduced adipogenesis, but only during late stages of differentiation [62]. These data suggest that SUMO may have temporal effects on adipogenesis. In support of this theory, we recently demonstrated that waves of sumoylation controls adipogenesis in 3T3-L1 cells by promoting the timely downregulation of preadipocyte genes and activation of mature adipocyte genes [33]. Moreover, in the present study, the pro-adipogenic effect of inhibiting SUMO was completely lost if the inhibitor was administered after day 2 of differentiation, and when administered only on days 4-9 it actually inhibited the differentiation in line with the abovementioned reports. Still, when the inhibitor was administered from day 0 and throughout differentiation, inhibition of sumoylation had a clear pro-adipogenic effect.

Other studies support our finding of SUMO as overall anti-adipogenic. For example, ablation of the desumoylases Senp1 or Senp2 has also been shown to inhibit adipogenesis via hypersumoylation of CEBPbeta, Setdb1 or Sharp1, resulting in impaired Pparγ activity [65–67]. Moreover, Pparγ itself can be sumoylated [68] and *in vitro* mutagenesis of the K107 sumoylation site clearly demonstrates the inhibitory effect of SUMO on Pparγ transcriptional activity in luciferase assays [69,70]. In the present study, we show a similar repressive effect of SUMO on Pparγ/Pgc-1α transcriptional activity. These findings could explain how downstream target genes of Pparγ were downregulated on day 1 of adipogenic differentiation in Irx3-KO cells while Pparγ itself was not. Moreover, while inhibition of sumoylation stimulated adipogenesis, we found it to repress osteogenesis in the ME3 *Irx3-*KO cells. Thus, our data suggest that sumoylation can promote osteogenesis. In agreement with this, Zhang et al. found TGFB-mediated sumoylation of Pparγ to inhibit adipogenesis and promote osteogenesis [31], and Liu et al. demonstrated a switch from osteogenic to adipogenic differentiation of bone-marrow-derived mesenchymal stem cells following pharmacological inhibition of sumoylation [30]. Taken together, our results extend our understanding by showing that Irx3 controls an adipocyte-osteocyte epigenetic switch in mesenchymal-like precursor cells, and this process is at least partially mediated via sumoylation.

Mesenchymal lineage determination is regulated by complexes of Trithorax group proteins, such as the COMPAS and SWI/SNF family members (as reviewed in [71,72]. The former includes various SET domain methyltransferases, including Setd1a, Setd1b and Kmt2a (Mll1) [71]. These genes were all found to be bound by Irx3 and upregulated following *Irx3-*KO (Figures 1F and 2D), and we previously found both Setd1a and Kmt2a to be sumoylated in adipocytes [33]. SET domain-specific truncation of Kmt2a in mice dramatically reduced mono-methylation of H3K4 on specific *Hox* genes, altering their expression levels and leading to skeletal defects [73]. Interestingly, SENP3-mediated desumoylation of Kmt2a complexes were found to be necessary for osteogenesis in dental follicle stem cells [74]. Thus, our data point to Setd1a and Kmt2a as possible mediators of the anti-osteogenic role of Irx3 in adipocyte precursor cells. However, their exact contribution remains to be determined.

While sumoylation clearly mediates a vital part of Irx3’s effect on adipogenesis, as demonstrated by the reversed phenotypes following ML-792 treatment, it likely does not mediate the entire effect. Most notably, we found Irx3 to bind the promoters and alter the expression of a broad range of histone modifying enzymes and chromatin remodelers, which may act independently of SUMO or at least constitute a different substrate pool for sumoylation. Secondly, while most of the SUMO-responsive genes were also changed by *Irx3-*KO, Irx3 additionally controlled a large set of genes that were unaffected by inhibition of sumoylation. For example, we identified Gas1 as a direct Irx3 target gene that increased 12-fold after *Irx3-*KO (Supplementary file 2). The Gas1 protein is a coreceptor that promotes Sonic Hedgehog (SHH) signaling, whose activation is marked by increased Gli1 expression (reviewed in [75]). Since we previously also found Gli1 expression to increase in response to *Irx3-*KO [14], our data clearly indicate increased SHH signaling in the *Irx3-*KO cells. The SHH pathway is well known to inhibit adipogenesis and promote osteogenesis [76,77], and is likely a contributing factor in the pro-osteogenic capacity of the *Irx3-*KO cells. The Gas1 protein was not found to be a SUMO substrate [33], thus the osteogenic effects of this factor is Irx3-dependent, but sumo-independent.

While our findings clearly implicate Irx3 in epigenetic regulation of adipogenic identity, extrapolating these results to explain a model where transient changes in IRX3 mediate the effect of *FTO* locus risk variants on human adipocyte beiging may not be accurate. First, our study was performed in mice and not humans. Secondly, while we have shown an epigenetic effect of complete Irx3 ablation, the epigenetic effect of more modest *FTO* locus-dependent alterations in IRX3 levels may be different. Thirdly, due to the profound effect on adipocyte differentiation *per se* obtained in this study, it was not possible to assess the effect on white versus beige development. Future studies should address these shortcomings.

### Conclusions

Our study uncovers Irx3 as a master transcriptional regulator of histone and chromatin remodeling enzymes in adipocyte precursor cells, promoting adipogenesis and suppressing osteogenesis. Moreover, we have identified Irx3 as the first reported upstream regulator of sumoylation, and found repression of sumoylation to be a key mediator of the pro-adipogenic effect of Irx3. These findings imply indirect epigenetic regulation of adipogenesis by Irx3 as a possible novel mechanism underlying the association between the *FTO* locus and obesity. More broadly, our study has identified a large set of novel direct IRX3 target genes, which may be relevant for understanding the role of IRX3 also in other conditions and diseases.

## Supporting information

Supplementary file 1_ChIP-seq

Supplementary file 2_ATAC-seq

Supplementary file 3_Direct_Irx3-target-genes_FC1.2

Supplementary file 4_SLAMseq_vs_Irx3-KO

Supplementary file 5_Quantseq_vs_Irx3-KO

Supplementary file 6_extended direct SUMO target genes and overlap with Irx3-KO

Supplementary file_7_ATAC-seq_Irx3-KO_vs_control

Supplementary file_8_GO ATAC-seq_prom Irx3-KO vs Ctr

Supplementary file_9_ATAC-seq vs RNAseq_Irx3-KO_D1_D7

## Acknowledgements

We thank the engineers at the Harwell Institute, Oxford, UK and the Hormone Laboratory at Haukeland university hospital, Bergen, Norway, particularly Michelle Simon, Carol Cook, Margit Solsvik and Linn Skartveit, for excellent technical assistance. We also thank Anagha Madhusudan Joshi-Michoel for valuable input in interpreting the ChIP-seq data. The WT ME3 cells were kind gifts from Karsten Kristiansen. The Genomics Core Facility (GCF) at the University of Bergen, which is a part of the NorSeq consortium, provided services on ATAC-sequencing.

## Author contributions

JIB, SL, MC, GM and SND conceived the study and designed the experiments with input from PC, KM, PRN, and RC. JIB performed the experiments pertaining to ME3 cells and analyzed the data from all experiments. SL performed the animal experiments and experiments pertaining to iWAT and gWAT. SY and NA performed the osteogenic and adipogenic experiments using 3D culture models. LD and XZ performed the bioinformatics. JIB made the figures and wrote the manuscript. All authors contributed to data interpretations, input on writing and approved the final version of the manuscript.

## Materials and methods

### Animals

Animal studies were conducted in accordance with the UK Animals (Scientific Procedures) Act (1986). Mouse studies were performed under the Home Office License 30/2642. Procedures were approved by the MRC Harwell Animal Welfare and Ethical Review Board (AWERB). Mice were housed according to UK Home Office welfare guidelines in a 12hr light / dark cycle at a temperature of 21 ± 2°C and humidity of 55 ± 10%. Mice were fed ad libitum and had free access to water (25 ppm chlorine). Mice that were used for primary pre-adipocyte isolation were fed SDS maintenance chow (RM3, 3.6 kcal/g).

### Cells

#### Isolation, culturing and differentiation of adipocyte precursor cells from gWAT and iWAT

Mouse primary adipocytes were isolated essentially as previously described [78]. Briefly, mouse WAT depots were excised from 6-10 weeks old C57BL/6NJ (B6N) mice and placed in PBS. Tissues were minced and digested in 5ml per depot digestion buffer (sterile Hank’s Balanced Salt Solution (HBSS; H8264), 0.8 mg/ml collagenase type 2 (Worthington Biochemical Corporation, NJ, USA; LS004174), 3% bovine serum albumin (BSA) (with fatty acids) and incubated in a 37°C water bath for 60-75 min with shaking every 10 min by hand. When digestion was complete, tubes were centrifuged for 3 min at 300xg to separate floating mature adipocytes from the supra-vascular fraction (SVF), containing adipocyte precursors. The supernatant containing the floating adipocyte fraction was removed and the cell pellet re-suspended in pre-warmed growth media consistent of DMEM GlutaMax (# 10569010 DMEM, high glucose, GlutaMAX™ Supplement, pyruvate) supplemented with 10% Fetal Bovine Serum (Gibco, New York, NY, USA 10082-147) and 1% Penicillin-Streptomycin (5,000 U/mL) (Gibco, 15070063) and grown at 37°C and 5% CO2. The cell solution was subsequently filtered through a 40 μm nylon mesh and plated on a 10 cm dish. Media was replaced the next day. For differentiation experiments, preadipocytes were seeded in density 100K cells/ml in 12-well or 6-well dishes and grown to confluence. 2 days post-confluence, preadipocytes were induced to differentiate using induction medium (DIFF1) containing growth medium supplemented with 0.5 mM 3-Isobutyl-1-Methylxathine (IBMX) (Sigma-Aldrich, St. Louis, MO, USA I5879), 1 μM Dexamethasone (Sigma-Aldrich, D2915), and 5 μg/ml human insulin (Sigma-Aldrich, I9278). At day 4 of differentiation, media was changed to maintenance media (DIFF2) containing growth medium supplemented with insulin only. Media was changed every 2 days and differentiation complete at day 7-9 post adipogenic induction. These cells were used for ATAC-seq and ChIP-seq as indicated.

#### ME3 WT and Irx3-KO cells

The wild type (WT) ME3 cell line was previously generated [79] and cultured and differentiated for adipogenesis as previously described [14]. Cells were harvested for ATAC-seq on days 0, 1 and 7 of differentiation in two independent experiments. CRISPR-Cas9 control and *Irx3-*KO cells were generated from WT ME3 cells, and their global gene expression was previously assessed by RNA-seq on days 1 and 7 of differentiation [14]. These cells were also used for adipogenic and osteogenic differentiation, qPCR, Bodipy staining, ATAC-seq, Western blotting (WB), Chromatin Immunoprecipitation (ChIP) and luciferase transactivation assays. For osteogenic differentiation, the cells were treated with 50 µg/mL L-ascorbic acid 2-phosphate, 10 nM dexamethasone and 10 mM β-glycerophosphate (all from Sigma-Aldrich-Aldrich) in the complete growth medium. The medium was changed every three days until day 14 for assessment of alkaline phosphatase (ALP) activity, or day 21 for Alizarin red S staining.

#### Cell differentiation in 3D

For adipogenic 3D differentiation, ME3 cells were first encapsulated in Gelatin methacryloyl (GelMA) hydrogel. Briefly, GelMA solutions were reconstituted by dissolving purified GelMA prepolymer powder (X-Pure GelMA 160P60 RG, Rousselot Biomedical, Belgium) in the growth medium in presence of 1 % w/v of Lithium Phenyl(2,4,6-trimethylbenzoyl)phosphinate (L0290, Tokyo Chemical Industries, Co., Japan) on a magnetic stirrer at 37 ℃ under sterile conditions in a dark environment. The ME3 cells were added in the solution with a density of 5 million cells per ml, which was then transferred onto molds in a diameter of 5 mm. Photo-polymerization was conducted using a dental curing lamp set for 1 single cycle of 10 seconds at a light intensity of 1200 mW/cm2. Adipogenesis was induced essentially as for 2D, using the same differentiation components as previously described [14], but the cells were kept two more days in the final medium for a total differentiation time of 9 days.

For osteogenic 3D differentiation, highly porous 3D scaffolds for osteogenic differentiation were fabricated as previously described [80]. Briefly, a co-polymer Poly(L-lactide-co-trimethylene carbonate) lactide was dissolved in chloroform and then mixed with sodium chloride particles ranging in size from 90 to 600 μm. After the complete evaporation of the solvent, the constructs were formed, each possessing a diameter of 12 mm and a thickness of 1.2 mm. To eliminate the presence of sodium chloride, the scaffolds were thoroughly washed with distilled water. The scaffolds were placed in 48 well plates where 500,000 cells in 300 μL of the culture medium were seeded per scaffolds. Osteogenic differentiation in 3D was performed as in 2D for 21 days as described above.

#### Lipid staining

Adipogenic differentiation was assessed on formaldehyde fixated cells by Oil-red-O lipid staining as previously described [14] or on live cells using Bodipy 493/503 (Sigma-Aldrich) lipid staining. For Bodipy staining, the culture medium was replaced with a Bodipy solution (1 µg/mL, freshly prepared in OptiMEM by vigorous vortexing) and incubated at dark at 37°C for 30 min. The staining solution was removed and replaced with regular growth medium before being imaged by fluorescent microscopy using a Leica SP8 confocal microscope (Leica, Germany) for 3D cultures and the Incucyte S3 Live-Cell Analysis System (Essen Biosciences, Ann Arbor, MI, USA) for 2D cultures.

#### Immunofluorescence

Samples from adipogenic and osteogenic 3D cultures were fixed in 4% PFA for 15 min followed by permeabilization in 0.1% Triton X-100 in PBS for 15 min at room temperature. The samples were then incubated in a blocking buffer consisting of 10% normal goat serum (NGS: ab7481; Abcam, USA) in 0.05% Tween-20 in PBS (PBSTw) for 60 min at room temperature. After blocking, the samples were incubated with anti-Runx2 antibody (1:250, ab23981; Abcam, USA) in PBSTw at 4°C overnight. After washing in PBSTw 6 times, the samples were incubated with goat anti-rabbit antibody Alexa Fluor 546 (1:500, A11010; Invitrogen) for 1 h at room temperature simultaneously with 4′,6-diamidino-2-phenylindole (DAPI: 1:2500, D9542; Sigma-Aldrich-Aldrich, USA) and Phalloidin Alexa488 (1:500, A12379; Invitrogen, USA) for nuclear staining and filamentous actin staining, respectively. Image acquisition was performed as z-stack images of 250 μm in depth in 3 separate channels using a Leica SP8 confocal microscope (Leica, Germany).

#### Alkaline phosphatase staining

Early osteogenic differentiation was assessed on day 14 and the collected samples were fixed with 4% paraformaldehyde (PFA) for a duration of 1 minutes at room temperature. Subsequently, the samples were incubated in BCIP®/NBT solution (B5655; Sigma-Aldrich-Aldrich) for 60 minutes at room temperature. For quantification, the substrate was extracted by incubating the samples with 100 mM cetylpyridium chloride overnight at room temperature. The absorbance of the extracted solution was measured at 540 nm using a Varioskan™ LUX multimode microplate reader (VLBL00D0; Thermo Scientific, Finland).

#### Alizarin red S staining

Late osteogenic differentiation was assessed on Day 21 and the collected samples underwent fixation in 4% PFA for 40 minutes. Subsequently, the samples were gently washed with Milli-Q® water. To visualize mineralized nodes, a 0.2% solution of Alizarin Red S (A5533; Sigma-Aldrich-Aldrich) was applied to the samples for 20 minutes, followed by six rinses with Milli-Q® water. For quantification, the dye was extracted from the samples using 100 mM cetylpyridium chloride, which was allowed to incubate overnight at room temperature. The dye extract was then subjected to absorbance measurement at 540 nm using the microplate reader.

#### Pharmacological inhibition of sumoylation

Cells were treated with 0.5 µM or 1 µM of the ML-792 sumoylation inhibitor on indicated days of adipogenic or osteogenic differentiation. The inhibitor specifically targets the E1 SUMO activating enzyme (Sae), leading to loss of global sumoylation without affecting neddylation or ubiquitination [81].

### ChIP-seq

#### Sample preparations

Primary pre-adipocytes from gWAT and iWAT of mice were isolated and differentiated as described above. For each depot, cells from 6-12 animals aged 41 days were pooled and digested. Cells were collected on days -1 and/or 1 as indicated in one to two independent experiments. Cells were fixated by addition of 1/10 volume of freshly prepared formaldehyde solution containing 11% formaldehyde, 0.1 M NaCl, 1mM EDTA pH 8.0 and 50 mM HEPES pH 7.9 to culture media. The fixation reaction was stopped after 15 minutes of agitation at RT by addition of 1/20 volume of 2.5 M glycine. The cells were thereafter kept on ice and washed twice with 0.5% Igepal in PBS. In the second wash, 1mM PMSF was added. Pellets were snap-frozen in liquid nitrogen and stored at -80°C until shipment on dry ice to Active Motif for ChIP-sequencing, where in-house protocols based on published methods were used [82–84].

#### ChIP-seq data analysis

Reads were aligned to mouse genome (mm10) using Bowtie2 (Version 2.3.4.3). Resulting filtered alignment files were first filtered to include only reads that were the primary alignment, with any duplicates and supplementary alignments being removed via samtools. Reads with a mapping quality below 30 were also removed. Filtered read sets ranged from 25 to 31.5 million reads.

Peak files were then annotated using the R package ChIPseeker [85] (Version 1.22.0) in order to annotate peak location in relation to known genomic features, as well as to generate maps of ChIP peak binding in reference to transcription start sites where peaks were within 3000 base pairs of a TSS.Log2 ChIP-over-input tracks for each alignment file were generated using deepTools bamCompare (Version 3.1.2), filtering out any reads from the Encode blacklist [Encode, mm10, 2014], and then uploaded to a UCSC track hub. Bam files were also submitted to Macs2 (Version 2.1.1) for peak calling, providing input data for each day as control. A q-value filter of 1e-4 was used to filter low-confidence peaks from the resulting Macs2 peak files. A total peak set of all bam files in the first replicate, as well as a merged signal graph track, were created for visualization.

ChIP-seq peaks were overlapped with peaks found in the ATAC-Seq data using the genomicRanges R package [86] (Version 1.38.0) and the enrichment of each overlapping peak, as derived from Macs2, was displayed via a heatmap. Simple correlation tests were performed to identify if the magnitude of enrichment for overlapping peaks was similar in both experiments. These data were then further overlapped with a previous *Irx3-*KO RNA-Seq dataset, generated as previously described [14] to highlight the time-dynamic effects of an Irx3 knockout on target regions which featured a gene promoter, an Irx3 ChIP-seq peak and a region of openness as defined by our ATAC-Seq experiment. Gene ontology enrichment analysis was performed on sets of peaks whose genomic location and annotation overlapped in different experiments via both the Panther [87] and the Reactome [88] pathway databases.

### ATAC-seq preparation

#### Preadipocytes from iWAT and gWAT

ATAC-seq was performed by adapting the protocol from [89]. Differentiating cells were lysed directly in cell culture plate. Lysis buffer was added directly onto cells grown in a 12-well plate. Plates were incubated on ice for 10 min until cells were permeabilized and the nuclei released. Lysis buffer was gently pipetted up and down to wash nuclei off the well and transferred into a chilled 1.5ml tube to create crude nuclei. Nuclei were spun down at 600 x g for 10 min at 4℃, nuclei pellets were then re-suspended in 40 µl Tagmentation DNA (TD) Buffer. Transposition reaction was performed as previously described [89]. All tagmented DNA was PCR amplified for 11-13 cycles. Quality was assessed using a DNA1000 Chip (Applied Biosystems) and run on a Bioanalyzer (Applied Biosystems). The profiles showed that all libraries had a mean fragment size of ∼190 bp and characteristic nucleosome patterning, indicating good quality of the libraries. Libraries were sequenced at the Wellcome Trust Centre for Human Genetics in Oxford on a HiSeq4000 Illumina generating 50 million reads/sample, 75bp paired end. To reduce bias due to PCR amplification of libraries, duplicate reads were removed.

#### WT ME3 cells

ME3 cells were collected for open chromatin profiling (ATAC-seq) by trypsination on days 0, 1 and 7 of differentiation in two independent experiments. For each sample, technical duplicates of 100 000 cells were frozen at a slow rate (-1°/minute) in 5% DMSO at -80°C in CoolCell freezing containers (BioCision, San Rafael, CA, USA). Open chromatin was profiled through a commercial service (Active Motif, Carlsbad, CA, USA) based on a method previously described [90]. In brief, chromatin was isolated from the cells followed by Tn5 transposase-mediated insertion of sequencing primers into open chromatin regions. After library preparation, high-throughput sequencing (Illumina) was performed and reads were then aligned to the mm10 genome using Bowtie2 (Version 2.3.4.3) using the “very-sensitive” setting.

#### Control and Irx3-KO ME3 cells

Cells were differentiated for 0, 1 and 7 days, pelleted and snap-frozen in liquid nitrogen. After thawing, cells were DNase treated and lysed in ice-cold ATAC seq lysis buffer from the ATAC-Seq Kit (Active Motif, Carlsbad, CA, USA) for 2 min. Crude nuclei extracts were spun down at 600xg for 10 min at 4°C and nuclei were resuspended in 40 µL 1X Tagmentation buffer. The integrity of the nuclei was verified under microscope, counted, and 50,000 nuclei was transferred pr sample for tagmentation. Tagmentation and indexing was performed according to the instructions of the ATAC-seq Kit (Active Motif). Library quality control was performed on a Bioanalyzer and sequenced at the Genomics Core Facility (GCF) at the University of Bergen using the HiSeq4000 Illumina and 75 bp paired end sequencing.

#### ATAC-seq Data Analysis

Reads were first filtered for mapping quality ≥ 30 (samtools Version 1.7-2) and duplicated reads, as well as reads mapping to the mitochondria, were subsequently removed (picardTools Version 2.8.1.1). Peak calling was performed via Macs2 (Version 2.1.1). Peak sets were first filtered using blacklisted regions from the ENCODE project [91], then converted into a simple annotation format for submission to the Rsubread featureCounts (Version 1.6.0) package. Using this consensus peak set as an annotation guide, each unmerged bam file was submitted to featureCounts and the subsequent count matrix was input into RStudio for peak annotation and visualization.

Time-course analysis for each peak was then performed via the R package maSigPro [92] (Version 1.56.0). A regression fit for each peak was performed, and the p-value associated with the F-statistic of each model was calculated to uncover peaks which change dependent on experimental group. Peaks with an adjusted p-value less than 0.1 were then selected, and stepwise regression was performed to select significant variables for each gene. Peaks which were found to change over time were then submitted to hierarchical clustering and grouped into six clusters with differing time-dependencies and visualized via ggplot2 using a glm smoothing method with polynomial regression.

To detect differential accessibility between control and *Irx3-*KO samples, individual peaks were first identified using Macs2 as described above, and sequencing reads within each individual peak were then counted using featureCounts [93] and DESeq2 [94].

#### RNA isolation, cDNA synthesis and real-time qPCR analysis

RNA was isolated either alone using the RNeasy mini kit (Qiagen, Hilden, Germany), the RNA/Protein Purification Plus kit (Norgen, Thorold, ON, Canada) or the Maxwell 16 LEV simplyRNA purification kit (Promega, Madison, WI, USA) according to the manufacturers’ instructions. RNA quality control, cDNA preparation and qPCR was performed as previously described [14], using the delta-delta Ct method relative to reference genes *Rps13* or *Hprt*. Primers were designed using either the Universal ProbeLibrary Assay Design Center (Roche, Basel, Switzerland) or Primer-BLAST softwares [95]. Primer sequences are shown in table 1.

**Table 1:**
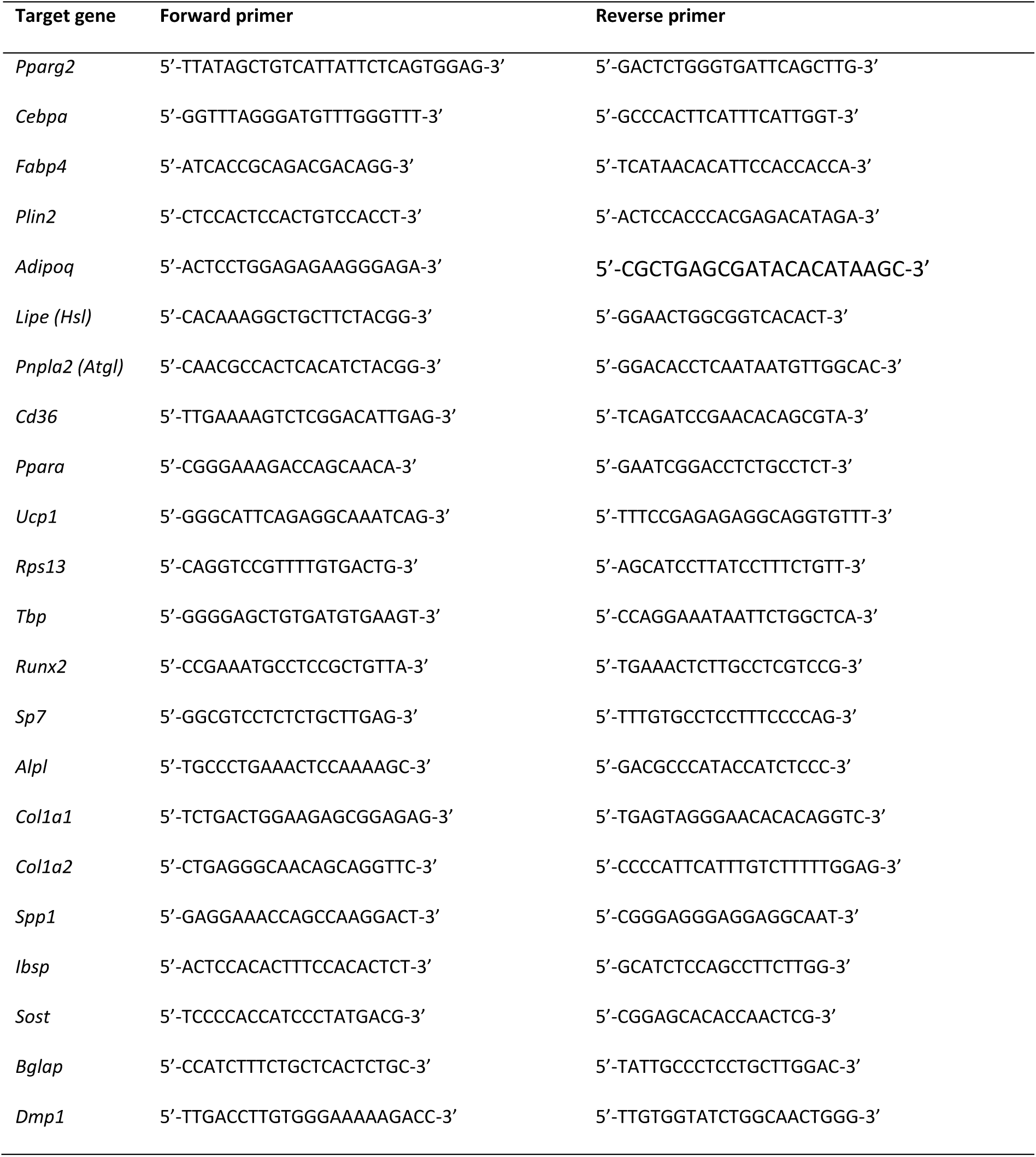
Primer sequences.

#### Protein isolation and Western Blotting

For isolation of total proteins with intact SUMOylome, fresh cells were washed in PBS and lysed directly in the culture plate with SUMO lysis buffer (2% SDS in PBS, pH 7.4, supplemented with 20 mM NEM and 2X cOmplete ULTRA protease inhibitor cocktail). Lysis was performed in SDS with NEM to immediately block and denature desumoylases, and at room temperature (RT) to avoid precipitation of SDS. Cells were sonicated at RT to reduce viscosity and boiled at 95°C for 5 min to facilitate protein unfolding and alkylation of free –SH groups through formation of stable thioether bonds. Samples were quantified using the SDS-compatible DC-kit (Biorad), followed by addition SDS-sample buffer containing excess DTT and boiled again at 95°C for 5 min to reduce disulphide bonds (does not affect the irreversible thioether bonds resulting from NEM treatment). All samples were analyzed by WB using 20 µg normalized lysate per sample. The following primary antibodies were used for WB: anti-Sumo2/3 (ab3742, Abcam), anti-AM (AB_2793779, Active Motif), anti-Actb (ab6276, Abcam) and anti-GAPDH (MAB374, Millipore). All primary antibodies were used at 1:1000 dilution. The following secondary antibodies were used: anti-mouse (554002, BD Biosciences) and anti-rabbit (31460, Invitrogen) at 1:7500 dilution.

#### Plasmids

The Irx3-AM construct was made by in-fusion cloning. Briefly, mouse Irx3 (NM_008393) was amplified by PCR from the donor vector pCMV6-entry-Irx3-myc-DDK (cat. no MR208149, Origene, Rockville, MD, USA) with the following forward 5’-*CGATCTAAGTAAGCTT*<u>CACC</u>ATGTCCTTCCCCCAGCTCG-3’ and reverse 5’-*GATCTTGGCAAAGCT*TAGACGAGGAGAGAGCTGATAAGACC-3’ primers. 5’ extensions (italic) matching HindIII digested pAM_1C empty destination vector (cat. no 53023, Active Motif) was added to each primer. A consensus Kozak sequence (underlined) was added to the forward primer just upstream of the start codon. The insert was cloned into the pAM_1C vector using the In-Fusion snap assembly master mix according to manufacturer’s instructions (Takara Bio, Kyoto, Shimogyō-ku, Japan). The purified plasmid was verified by Sanger sequencing.

#### Luciferase reporter assays

ME3 cells were seeded in 96-well plates at 7500 cells/well and grown for 24h before transfected with 20 ng of the pGL3-TK-PPRE-firefly-luc reporter construct, 1 ng of the constitutively expressed pRL-SV40-Renilla normalizator construct and 40 ng each of pCDNA-Pparg2 and/or pCDNA-Pgc-1a. The pCDNA-empty vector was used to keep total the total plasmid levels constant at 101 ng/well. The plasmids were diluted in OptiMEM and cells were transfected using TransIT-LT1 at a DNA:reagent ratio of 1:5. One day after transfection, the culture medium was replaced with fresh medium containing either DMSO or 0.5 µM ML-792 (MedChemExpress, Monmouth Junction, NJ, USA) in combination with either DMSO or 10 µM rosi. Two days after transfection, the cells were lysed and analyzed using the twinlite kit (PerkinElmer, Waltham, MA, USA) and the FLUOstar OPTIMA luminometer (BMG Labtech, Ortenberg, Germany) according to manufacturers’ instructions.

#### Statistical analysis

Omics data were analyzed in R as described above. Remaining data was analyzed and graphed using GraphPad Prism (9.2.0) and is shown as mean ± SD. Statistically significant differences between group means were investigated using Student’s unpaired t-test, one-way ANOVA or two-way ANOVA with Holm-Sidak correction for multiple testing as indicated. Data were tested for normality and homogeneity of variance and log_e_-transformed if necessary prior to statistical analysis.

## Supplementary data

### Supplementary figures

**Supplementary Figure S1:**
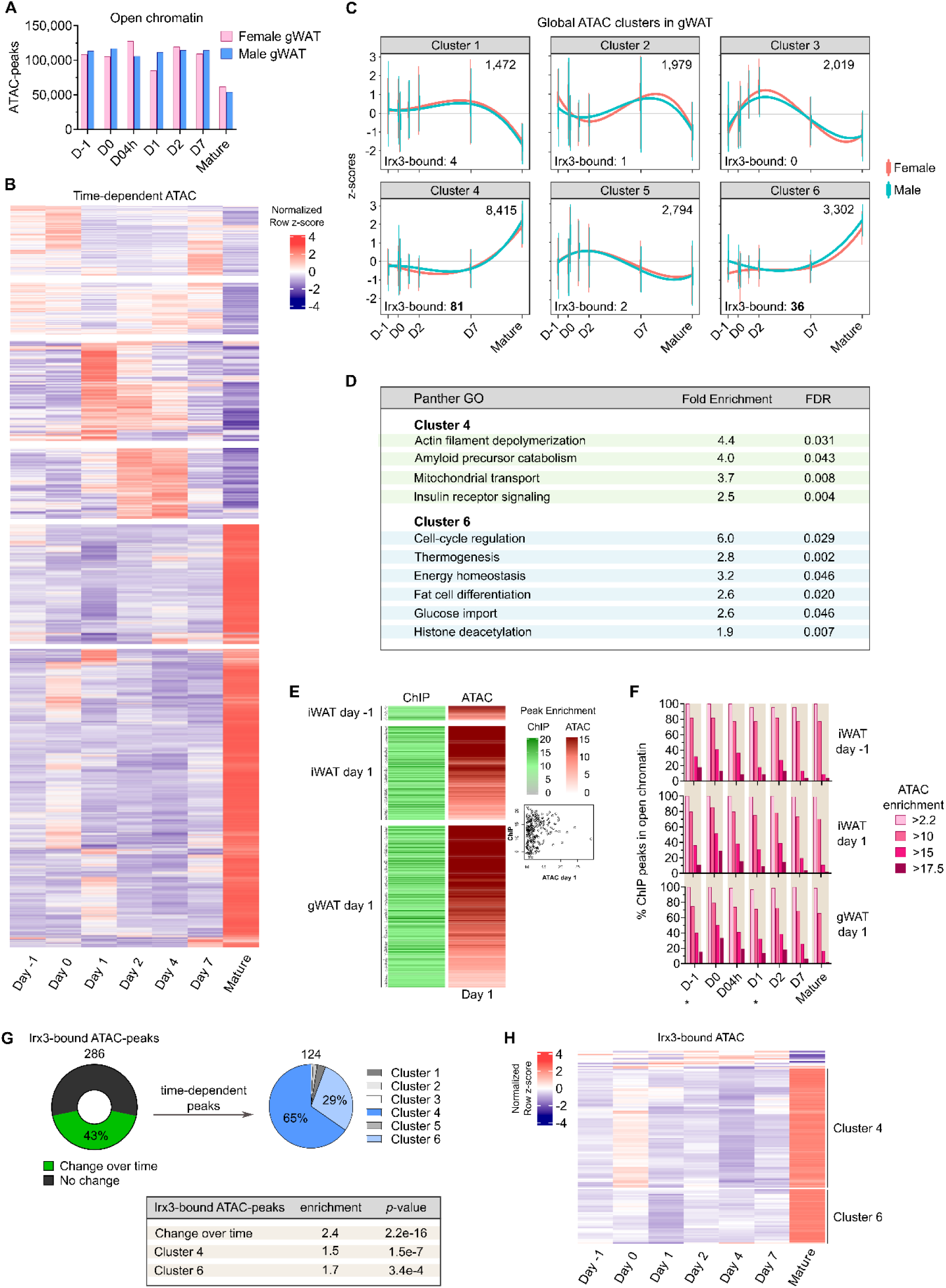
Irx3 binds to increasingly open chromatin in gWAT. Open chromatin profiling was performed by ATAC-sequencing at seven timepoints during differentiation of preadipocytes derived from gWAT of male and female mice. A) Total number of ATAC peaks shown for each timepoint during gWAT differentiation. B) Heatmap of the ATAC peaks that displayed significant time-dependent change in magnitude. Normalized row z-scores shown. C) Clustering of time-dependent ATAC peaks according to maSigPro stepwise regression. The total number of ATAC peaks, as well as the number of Irx3-bound ATAC peaks shown. D) Selected Panther GOs of the two clusters comprising loci with increased chromatin opening during differentiation. Fold enrichment and FDR shown. See also Supplementary file 2 for all GOs. E) Heatmap comparing enrichment of ChIP-seq peaks from all conditions (iWAT and gWAT) with ATAC-seq peaks in gWAT day 1. F) Percentage of ChIP-seq peaks overlapping ATAC-seq peaks for each timepoint during differentiation. Asterisks denote the timepoints where ChIP-seq was performed and thereby are directly comparable with ATAC-status. Color of bars represent degree of ATAC-enrichment/chromatin openness. G) Percentage of Irx3-bound ATAC peaks displaying changes over time (left). Distribution of Irx3-bound, time-dependent ATAC peaks according to the six global ATAC clusters (right). Enrichment of time-dependent Irx3-bound versus total ATAC-seq peaks shown (bottom). H) Heatmap illustrating the high proportion of Irx3-bound, time-dependent ATAC peaks that belong to cluster 4 and 6; loci that display increased opening in mature adipocytes. Normalized row z-scores shown. See (B) for corresponding heatmap of total ATAC peaks

**Supplementary Figure S2:**
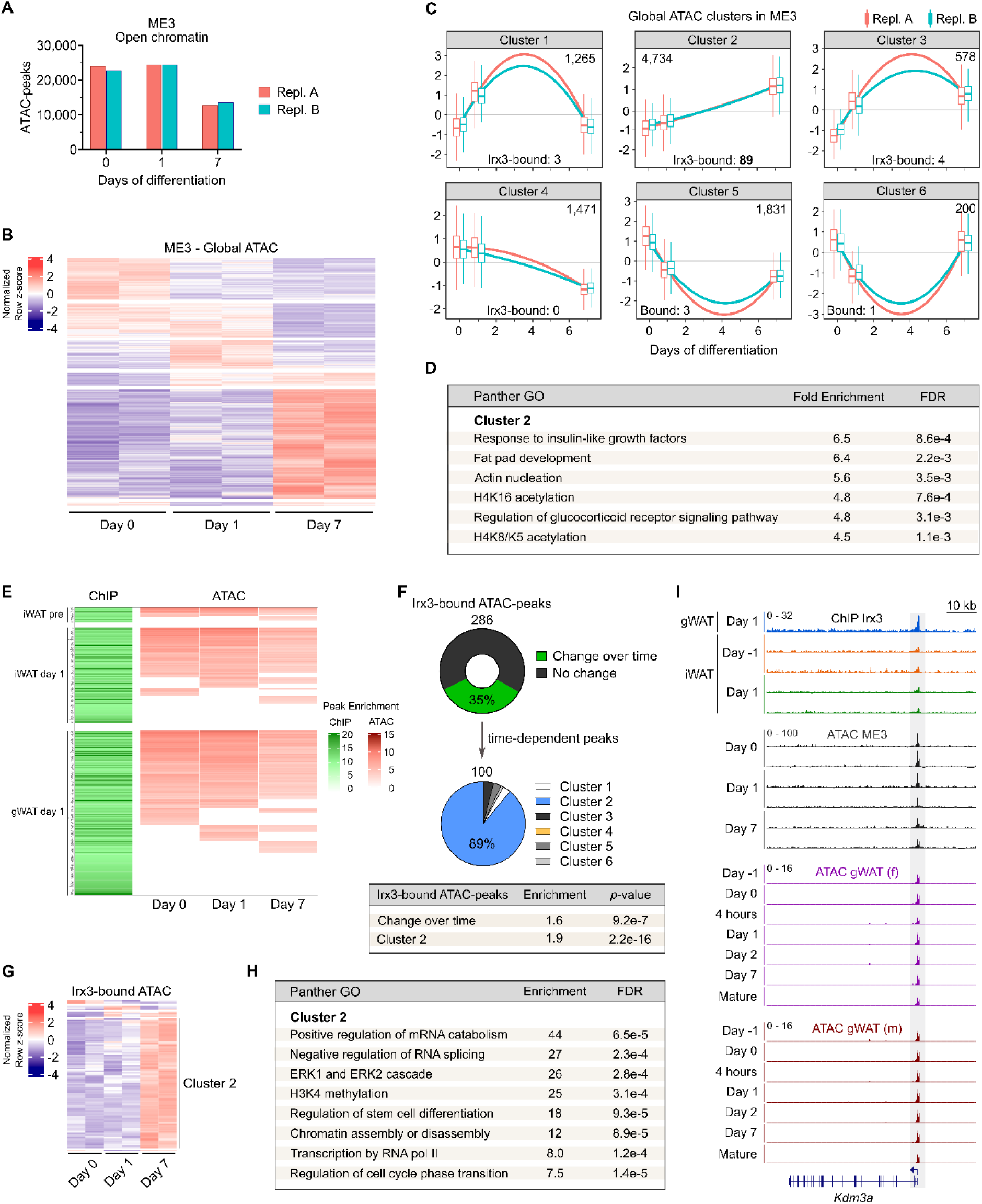
Irx3 binds to increasingly open chromatin in beige ME3 cells. Open chromatin profiling was performed by ATAC-sequencing.at three timepoints during differentiation of beige ME3 preadipocytes A) Total number of ATAC peaks shown for each timepoint during ME3 differentiation. B) Heatmap of the ATAC peaks that displayed significant time-dependent change in magnitude. Normalized row z-scores shown. C) Clustering of time-dependent ATAC peaks in ME3 according to maSigPro stepwise regression. The total number of ATAC peaks, as well as the number of Irx3-bound ATAC peaks shown. D) Selected Panther GOs of cluster 2, comprising loci with increased chromatin opening during differentiation. Fold enrichment and FDR shown. See also Supplementary file 2 for all GOs. E) Heatmap comparing enrichment of ChIP-seq peaks from all conditions (iWAT and gWAT) with ATAC-seq peaks in ME3 days 0, 1 and 7. F) Percentage of Irx3-bound ME3 ATAC peaks displaying changes over time (top). Distribution of Irx3-bound, time-dependent ATAC peaks in the six global ME3 ATAC clusters (middle). Enrichment of time-dependent Irx3-bound versus total ATAC-seq peaks shown (bottom). G) Heatmap illustrating the high proportion of Irx3-bound, time-dependent ME3 ATAC peaks that belong to cluster 2; loci that display increased opening in mature adipocytes. See (B) for corresponding heatmap of total ME3 ATAC peaks. H) Panther GOs for genes bound by Irx3 and displaying increased chromatin opening during ME3 differentiation. I) Genome browser aligning ChIP-seq tracks with ATAC-seq tracks from all timepoints in ME3 cells, gWAT cells on the promoter of the Kdm3a gene.

**Supplementary Figure S3:**
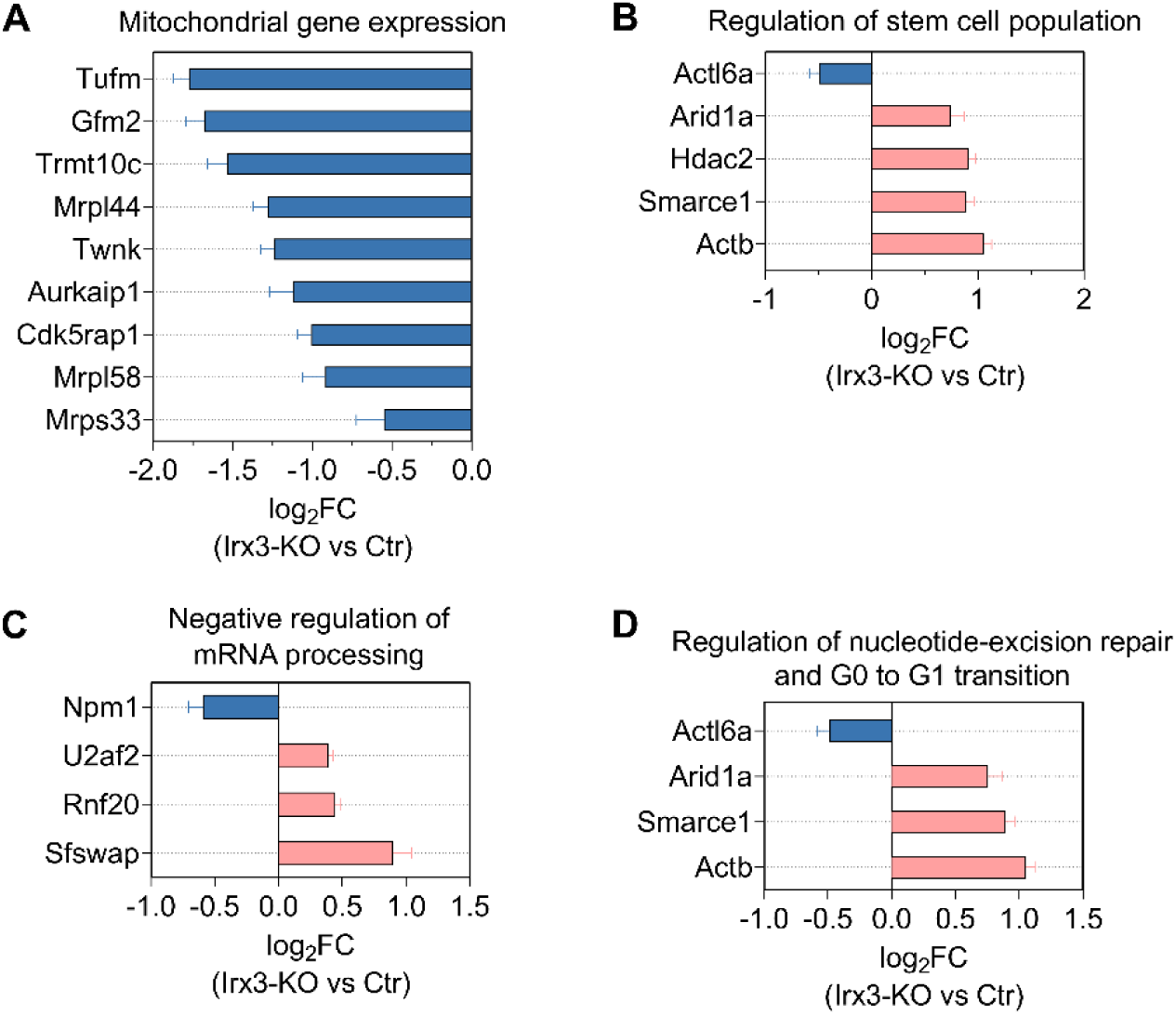
Expression of direct Irx3 target genes in *Irx3*-KO cells from indicated GO categories on day 7 of differentiation. Log2-fold changes for each gene in *Irx3*-KO vs control cells for each GO category is shown. Downregulated genes, blue bars; upregulated genes, red bars. Related to Figure 2.

**Supplementary Figure S5:**
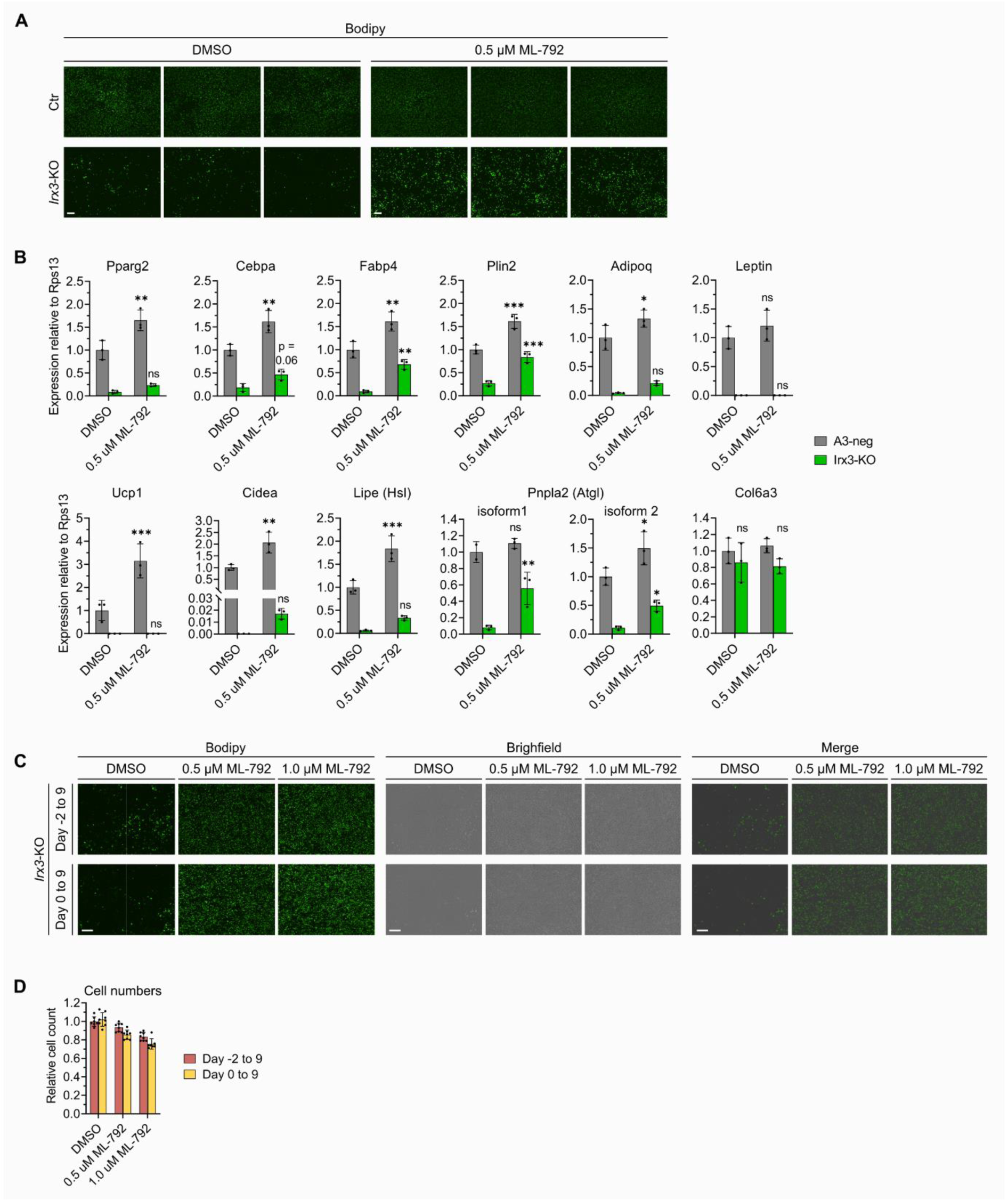
Inhibition of sumoylation partially restores adipogenesis in *Irx3-*KO cells. A) ME3 control and *Irx3-*KO cells were treated with either vehicle or the SUMOylation inhibitor ML-792 from days -2 to 9 of adipogenic differentiation. On day 9, the cells were treated with the Bodipy fluorescent lipid stain for 1 hour and imaged with fluorescence microscopy. Nine images/well were stitched to represent each well of a 24-well plate. Bodipy signal in triplicate wells from one of four independent experiments is shown. Scale bar = 400 µm. B) Gene expression in ME3 control and *Irx3-*KO cells in response to 0.5 µM ML-792, relative to *Rps13*. *p < 0.05, **p < 0.01, ***p < 0.001, two-way ANOVA with Holm-Sidak correction for multiple testing. C) Dose-response effect of ML-792 on lipid accumulation in *Irx3-*KO cells when ML-792 was added on either day -2 or 0 and kept throughout differentiation until day 9. Four images from each well of a 96-well plate were stitched together. One representative well from one of three independent experiments is shown with Bodipy fluorescent signal (left), brightfield (middle) and merge (right). D) Dose-response effect of ML-792 on relative cell count in *Irx3-*KO cells. n = 8 replicate wells from one of three independent experiments. Related to Figure 4.

**Supplementary Figure S6:**
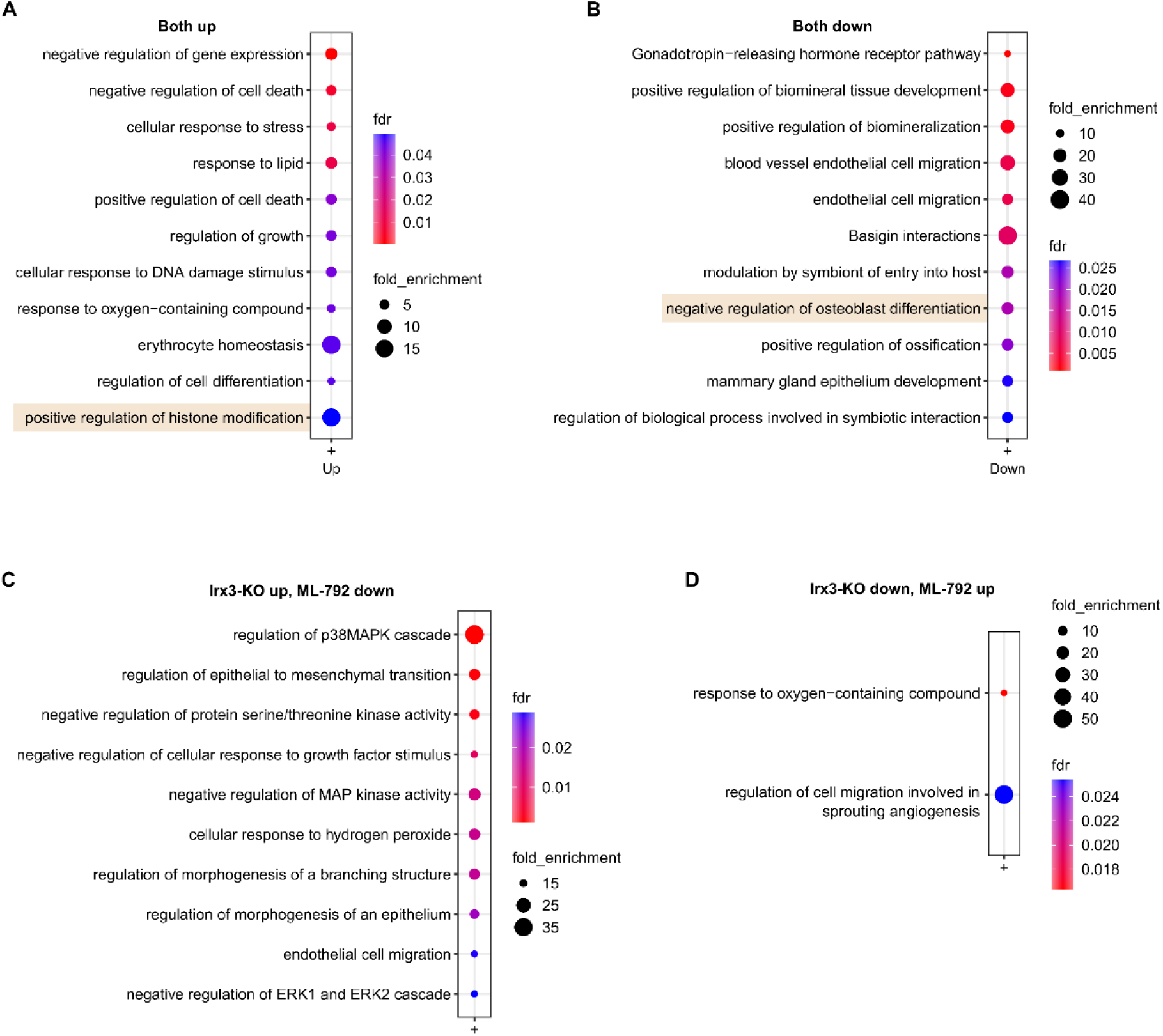
GOs for genes responding to both *Irx3-*KO (RNA-seq) and ML-792 (SLAM-seq) on day 1 of adipogenesis. Global mRNA-sequencing was performed to detect differentially expressed genes (DEGs) between ME3 control and *Irx3-*KO cells on day 1 of differentiation, using FC ≥1.2 and padj. ≤0.001. SLAM-seq was performed to detect changes in newly transcribed genes between 3T3-L1 cells treated with vehicle or 0.5 µM ML-792 on day 1 of differentiation, using FC ≥1.2 and padj. ≤0.05. Overlapping DEGs were subjected to combined GO analyses from Panther Pathways, Panther Biological processes complete and Reactome pathways. Top enriched/most significant categories are shown. Related to Figure 6. For complete lists, see Supplementary file 4. A) Top GOs for upregulated genes in both datasets. B) Top GOs for downregulated genes in both datasets. C) Top GOs for genes up- and downregulated with *Irx3-*KO and ML-792, respectively. D) Top GOs for genes down- and upregulated with *Irx3-*KO and ML-792, respectively.

**Supplementary Figure S7:**
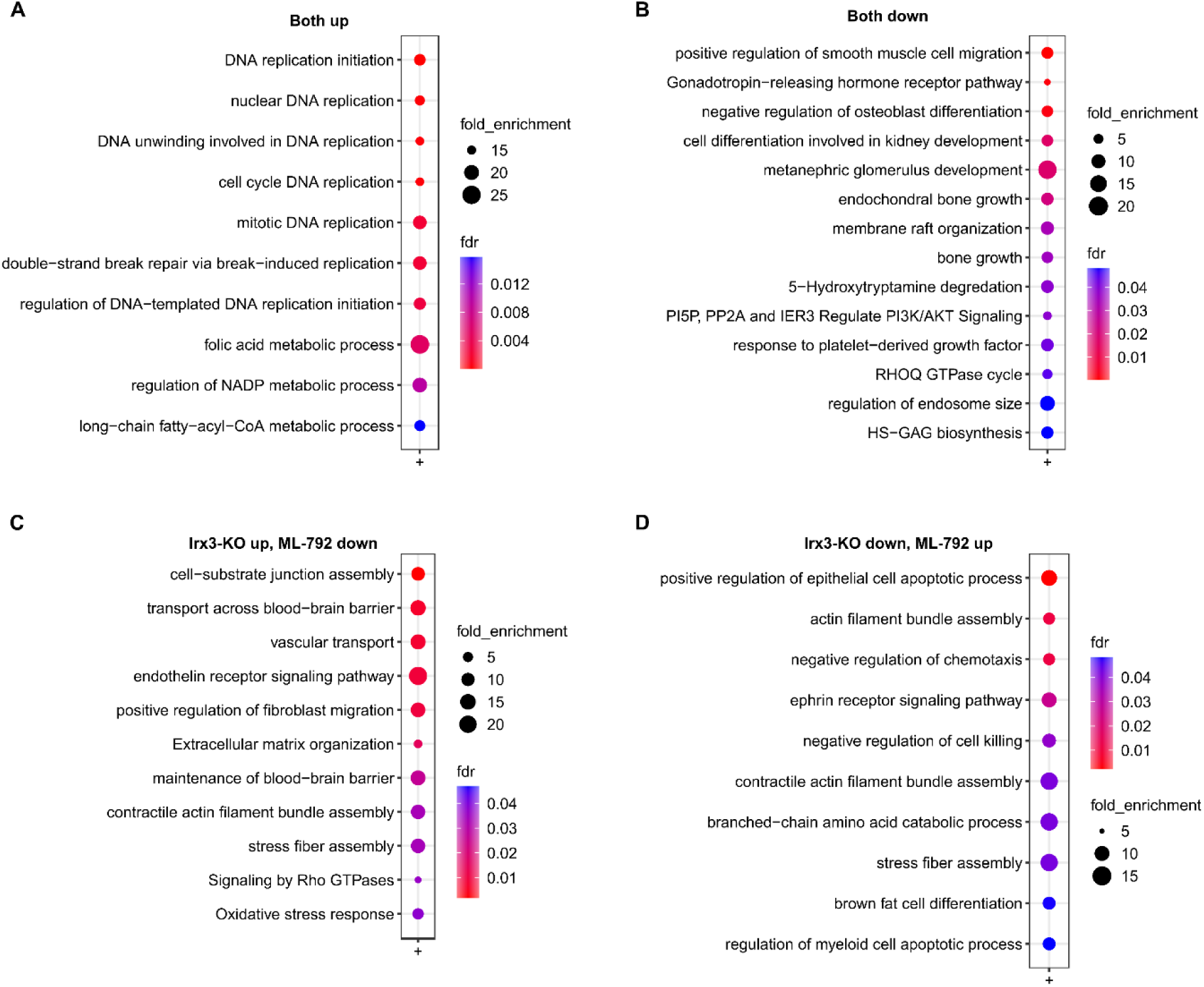
GOs for genes responding to both *Irx3-*KO and ML-792 (RNA-seq) on day 1 of adipogenesis. Global mRNA-sequencing was performed to detect differentially expressed genes (DEGs) between ME3 control and *Irx3-*KO cells on day 1 of differentiation, using FC ≥1.2 and padj. ≤0.001 and DEGs between 3T3-L1 cells treated with vehicle or 0.5 µM ML-792 on day 1 of differentiation, using FC ≥1.2 and padj. ≤0.001. Overlapping DEGs were subjected to combined GO analyses from Panther Pathways, Panther Biological processes complete and Reactome pathways. Top enriched/most significant categories are shown. Related to Figure 6. For complete lists, see Supplementary file 5 A) Top GOs for upregulated genes in both datasets. B) Top GOs for downregulated genes in both datasets. C) Top GOs for genes up- and downregulated with *Irx3-*KO and ML-792, respectively. D) Top GOs for genes down- and upregulated with *Irx3-*KO and ML-792, respectively.

**Supplementary Figure S8:**
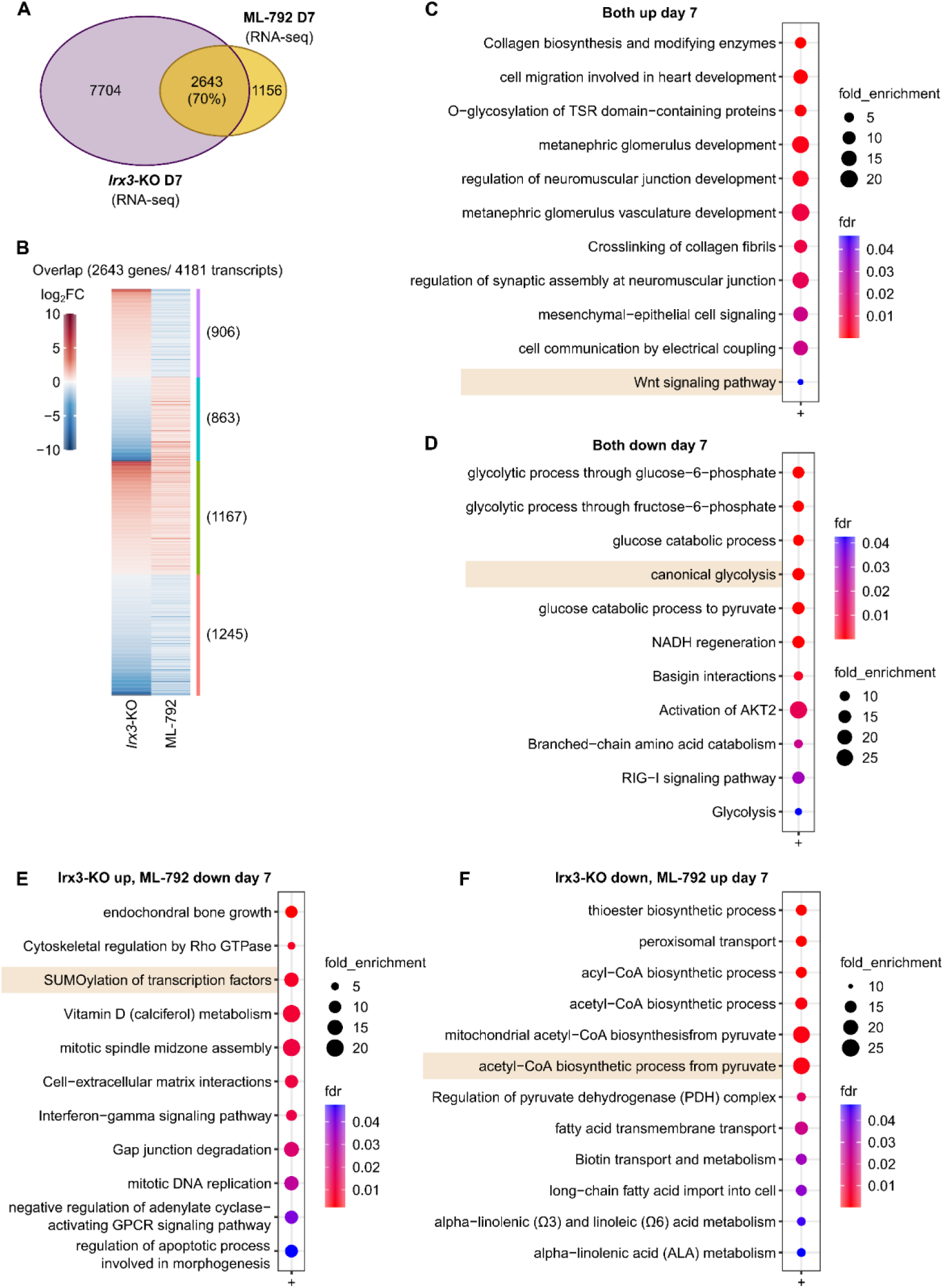
Overlap between Irx3 and sumo target genes on day 7 of adipogenesis. Global mRNA-sequencing was performed to detect differentially expressed genes (DEGs) between ME3 control and *Irx3-*KO cells on day 7 of differentiation, using FC ≥1.2 and padj. ≤0.001 and DEGs between 3T3-L1 cells treated with vehicle or 0.5 µM ML-792 on day 7 of differentiation, using FC ≥1.2 and padj. ≤0.001. A) Venn diagram showing overlap between DEGs following *Irx3-*KO or ML-792 on day 7 of differentiation. B) Heatmap showing direction of change among overlapping genes in (A). C) Selection of top enriched/most significant GO categories among the overlapping genes in (A-B).

**Supplementary Figure S9:**
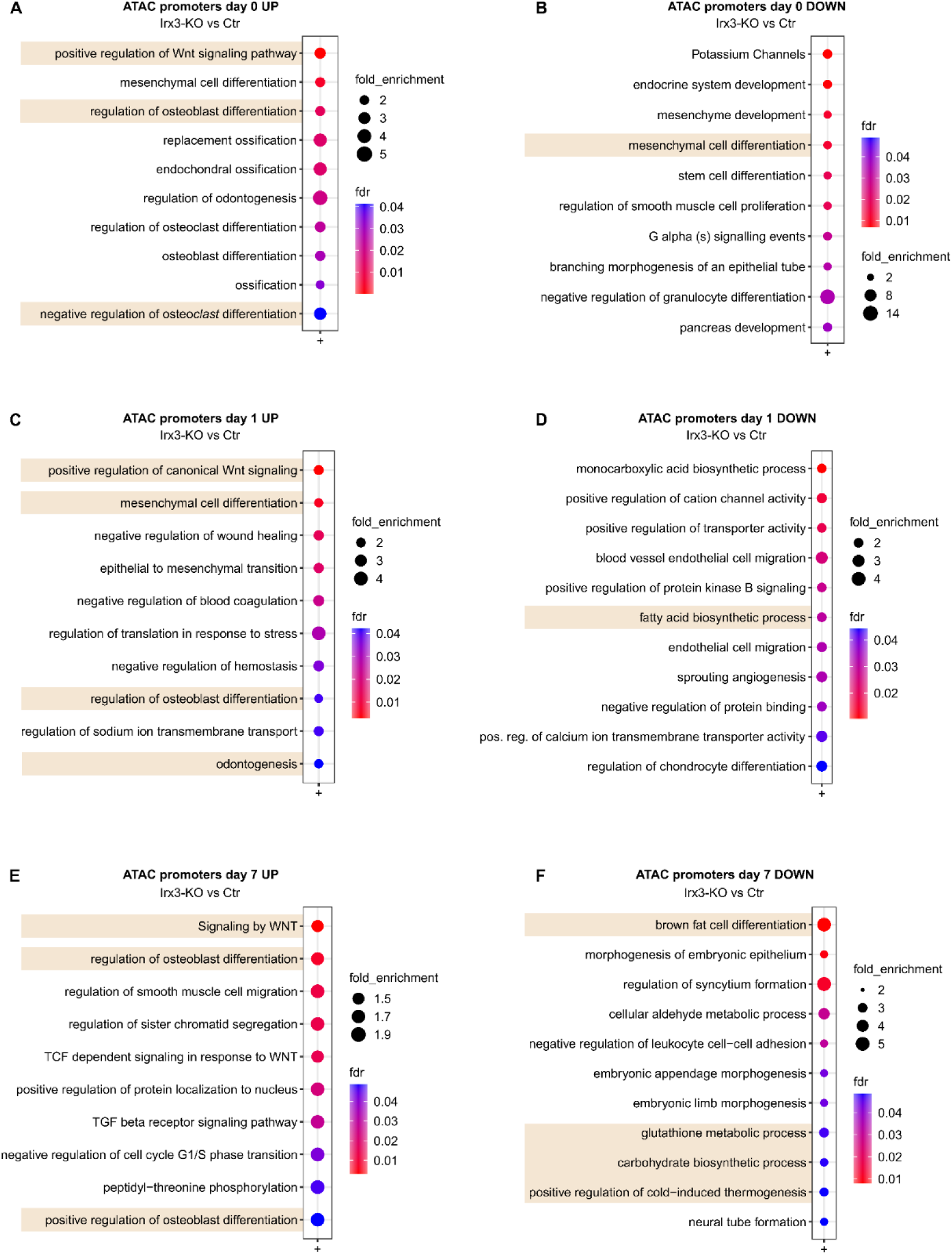
GOs for differential ATAC-seq peaks in promoters of *Irx3-*KO vs control cells. ME3 control and *Irx3-*KO cells were treated with adipogenic stimuli for 7 days, and open chromatin profiling on promoters was performed by ATAC-seq on days 0, 1 and 7. Most enriched and/or significant GOs for more open chromatin on day 0 (A), day 1 (C) and day 7 (E) and less open chromatin on day 0 (B), day 1 (D) and day 7 (F) shown. See supplementary file 8 for complete lists.

**Supplementary Figure S10:**
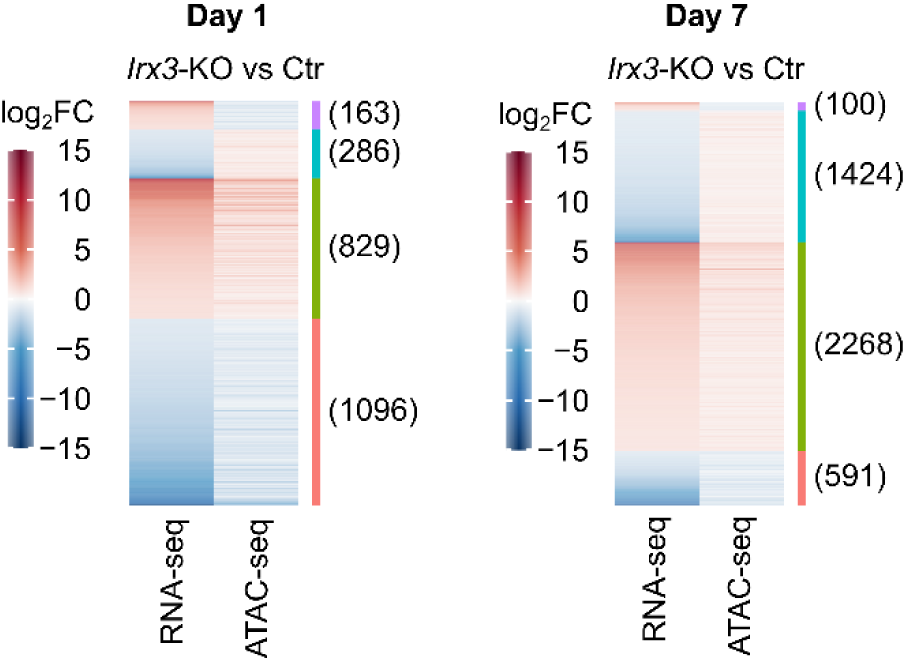
Changes in open chromatin and gene expression following *Irx3-*KO. Global changes in open chromatin in promoters of *Irx3-*KO vs control ME3 cells was measured by ATAC-seq and compared with changes in gene expression by RNA-seq. Log2-fold changes on day 1 (left) and day 7 (right) is shown.

